# Phosphoregulation of HORMA domain protein HIM-3 promotes asymmetric synaptonemal complex disassembly in meiotic prophase in *C. elegans*

**DOI:** 10.1101/2020.07.01.182063

**Authors:** Aya Sato-Carlton, Chihiro Nakamura-Tabuchi, Xuan Li, Hendrik Boog, Madison K Lehmer, Scott C Rosenberg, Consuelo Barroso, Enrique Martinez-Perez, Kevin D Corbett, Peter Mark Carlton

## Abstract

In the two cell divisions of meiosis, diploid genomes are reduced into complementary haploid sets through the discrete, two-step removal of chromosome cohesion, a task carried out in most eukaryotes by protecting cohesion at the centromere until the second division. In eukaryotes without defined centromeres, however, alternative strategies have been innovated. The best-understood of these is that used by the nematode *Caenorhabditis elegans*, where upon division of the chromosome into two segments or arms by the single off-center crossover, several chromosome-associated proteins or post-translational modifications become specifically partitioned to either the short or long arm, where they affect the timing of cohesion loss through as-yet unknown mechanisms. Here, we investigate the meiotic axis HORMA-domain protein HIM-3 and show that it becomes phosphorylated at its C-terminus, within the conserved “closure motif” region bound by the related HORMA-domain proteins HTP-1 and HTP-2. Binding of HTP-2 is abrogated by phosphorylation of the closure motif in *in vitro* assays, strongly suggesting that *in vivo* phosphorylation of HIM-3 likely modulates the hierarchical structure of the chromosome axis. Phosphorylation of HIM-3 only occurs on synapsed chromosomes, and similarly to previously-described phosphorylated proteins of the synaptonemal complex, becomes restricted to the short arm after designation of crossover recombination sites. Regulation of HIM-3 phosphorylation status is required for timely disassembly of synaptonemal complex central elements from the long arm, and is also required for proper timing of HTP-1 and HTP-2 dissociation from the short arm. Phosphorylation of HIM-3 thus plays a role in establishing the identity of short and long arms, thereby contributing to the robustness of the two-step chromosome segregation.

## Introduction

Meiosis reduces chromosome number from diploid to haploid by carrying out two rounds of sequential chromosome segregation following a single round of DNA replication. To segregate chromosomes correctly over the course of two divisions, two distinct chromosomal domains must be established as sites of cohesion loss for the first and second meiotic divisions, where cohesin complexes joining sister chromatids at each site are degraded stepwise. Organisms whose chromosomes have single, defined centromeres (monocentric) lose cohesin from chromosome arms in meiosis I, while cohesin at centromeres is protected by the protein Shugoshin [reviewed in l] until meiosis II. In contrast, organisms with holocentric chromosomes, such as *Caenorhabditis elegans*, must define these two separation domains with a different mechanism. In *C. elegans*, each chromosome receives one and only one crossover (CO) [reviewed in 2], which divides the chromosome into two unequal domains termed the short and long arms [3,4]. At meiosis I, cohesion is removed specifically on short arms, while cohesion on long arms persists until is finally degraded at meiosis II, thus separating chromatids in two discrete steps [reviewed in 5]. Although crossover position is biased towards the terminal thirds of chromosomes [6], crossovers can potentially occur at any position along the chromosome. Therefore, establishment of short and long arm domains must occur facultatively for each chromosome in each meiocyte.

The synaptonemal complex (SC) is a protein macroassembly that plays a critical role in holding homologous chromosomes together to ensure formation of crossovers during meiotic prophase [reviewed in 7]. Once crossovers are formed, the SC is disassembled in a stepwise manner, which influences the timing of cohesin degradation during the two subsequent meiotic divisions. In both monocentric and holocentric organisms, SC components disassemble in a non-uniform manner [4,8-13]. SC components are actively maintained at paired centromeres, and disassemble from chromosome arms, in diplotene and diakinesis in budding yeast, flies and mice, while in contrast, specific SC components are maintained either at short or long arms after SC disassembly in worms. This stepwise or asymmetric disassembly of SC components is suggested to promote centromere bi-orientation and kinetochore function, and establish chromosome separation domains [14]. However, the mechanisms regulating asymmetric SC disassembly are poorly understood.

In *C. elegans*, multiple upstream factors interacting with the SC, as well as post-translational regulation of SC components, have been found to be important for chromosome partitioning into short and long arms. The 3 ligases ZHP-1 and ZHP-2, that act with partners ZHP-3 and ZHP-4 to ensure the formation of a single crossover per chromosome, localize to short arms and are required for asymmetric SC disassembly [15]. In addition, phosphoregulation of the SC transverse and central elements, SYP-1 and SYP-2, is known to promote establishment of short and long arms [16,17]. MAP kinase-mediated phosphorylation of SYP-2 and subsequent inactivation of MAPK upon crossover designation triggers disassembly of SC central element proteins from the long arm [17]. Phosphorylation at the Polo Box Domain (PBD) binding motif of SYP-1 at Thr452 promotes the cascade of short and long arm establishment by recruiting PLK-2 to the SC. Phosphorylated SYP-1 and PLK-2 are mutually dependent for their colocalization on short arms upon crossover designation, and a non-phosphorylatable *syp-1* mutation leads to loss of asymmetry and failure in meiosis I segregation [16]. Once short/long asymmetry is established, the chromosome passenger complex (CPC) containing INC NP^ICP-1^ and Aurora B^AIR-2^ is recruited to the short arm by Haspin kinase phosphorylation of histone H3Thr3 [16,18-20]. H3Thr3 in turn is dephosphorylated on the long arm by HTP-1/LAB-1-based recruitment of phosphatase PP1^GSP-2^ [18,21]. AuroraB^AIR-2^ then phosphorylates cohesin R C-8 on the short arm, allowing its removal at meiosis I [18-20]. Taken together, these studies have shown that asymmetric partitioning of SC components and their interactors collectively ensures the establishment of chromosome domains required for correct segregation. However, the mechanisms that sense the crossover intermediates, partition chromosomes in a length-sensitive manner and disassemble SC components in an asymmetric manner are not understood well.

The HORMA domain family proteins (HORMADs) were first identified by sequence conservation among <u>Ho</u>p1, <u>R</u>ev7 and <u>Mad</u>2 in budding yeast, and act in cellular processes such as meiosis, mitotic cell cycle control and the spindle assembly checkpoint [22,reviewed in 23]. Previous studies have shown that the HORMA domain contains a C-terminal region termed a “safety belt” which wraps around, and locks in place, the “closure motif” coil of a binding partner [24].There are four HORMA domain proteins that make up the meiotic chromosome axis in *C. elegans*: the paralogs HTP-1 and HTP-2, which share 82% sequence identity and have nearly identical HORMA domain structures, HTP-3, and HIM-3 [24-28]. Each protein contains an N-terminal HORMA domain and at least one C-terminal closure motif [24]. With HTP-3 as the platform of assembly, these four proteins assemble in a hierarchical manner to form the axis, in which HTP-1/2 bind to HIM-3’s sole closure motif as well as HTP-3’s closure motifs 1 and 6, while HIM-3 binds to HTP-3’s closure motifs 2, 3, 4 and 5 [24]. Once crossover intermediates are formed, HTP-1/2 are removed from short arms and persist on long arms to recruit LAB-1 and PP1^GSP-2^ for protection of cohesion on long arms, while HIM-3 and HTP-3 remain on both short and long arms [3,18,21,29]. The molecular mechanisms removing HTP-1/2 from short arms, and SC central element proteins from long arms, in response to crossover designation remains a mystery.

Here, we characterize phosphorylation sites within the closure motif of HIM-3, and find that this phosphorylation promotes asymmetric SC disassembly as meiocytes exit pachytene in *C. elegans*. Phosphorylated HIM-3 localizes along the entire length of the SC in early meiotic prophase, and later becomes enriched at short arms once crossover intermediates are formed. We found that under conditions with limited crossover formation, this asymmetric partitioning of phosphorylated HIM-3 fails when fewer than four crossovers are present, suggesting that a dosage-sensitive mechanism triggers chromosome partitioning. Mutations in *him-3* phosphoresidues also delay asymmetric disassembly of the SC upon the exit from pachytene, suggesting that phosphoregulation of HIM-3 is part of the multilayered process establishing chromosome arm identity and thus ensuring meiotic chromosome segregation.

## Results

### HIM-3^HORMAD1/2^ is phosphorylated at the conserved closure motif

To understand regulation of meiotic proteins by post-translational modification, we previously carried out mass spectrometry to identify phosphorylated proteins in *C. elegans* adult hermaphrodites [16]. In two independent experiments, we found that the chromosome axis protein HIM-3 is phosphorylated at its C-terminus. We detected phosphopeptides containing Ser282 either singly or in combination with Ser274, Ser277, or Tyr279, and the most commonly-observed peptide contained doubly-phosphorylated Ser277/Ser282 (**Figure 1A** and **Supplemental Table 1**). Among the three most C-terminal phosphoresidues, which reside within the conserved closure motif, Tyr279 and Ser or Thr at position 282 are conserved features among nematode HIM-3 orthologs (**Figure 1A**). Mouse HORMAD1 Ser375 was shown to be phosphorylated on unsynapsed chromosomes, and this phosphorylation is hypothesized to signal the status of synapsis [30]. More recent studies further identified this C-terminus region containing Ser375 to be the closure motif, able to be bound by the HORMA domain of HORMAD1 itself and HORMAD2 [24,31]. While phosphorylation at the closure motif is conserved between *C. elegans* HIM-3 and mouse HORMAD1 (**Supplemental Figure 1A**), the function is not understood. Since other HORMA family proteins, HTP-1 and −2, are known to bind the HIM-3 closure motif in *C. elegans* [24], we first examined if phosphorylation affects HTP-1/2 binding to HIM-3 *in vitro*. A previous study has shown that both HTP-1 and −2 proteins bind unmodified HIM-3 peptides containing the closure motif in a fluorescence polarization (FP) peptide binding assay [24]. Examination of the crystal structure resolved by the same study led us to predict that phosphorylation at Ser282 would likely interfere with HTP-1/2 binding (**Figure 1B**). We conducted a similar FP peptide binding assay using purified HTP-2 and fluorescently tagged HIM-3 closure motif peptides containing various modifications (**Figure 1C**). As expected, HTP-2 proteins bound to the unmodified HIM-3 peptide (*Kd*=5.8µM). In contrast, phosphorylation at Ser282 either solely or in combination with phosphorylation of other residues completely abrogated HTP-2 binding to HIM-3 closure motif peptides (*Kd*: no significant binding detected), while phosphorylation at HIM-3 Tyr279 mildly lowered binding (*Kd*=16.1µM). This suggested the possibility that HTP-1/2 binding to HIM-3 is modulated by HIM-3 phosphorylation status at the closure motif.

**Figure 1.**
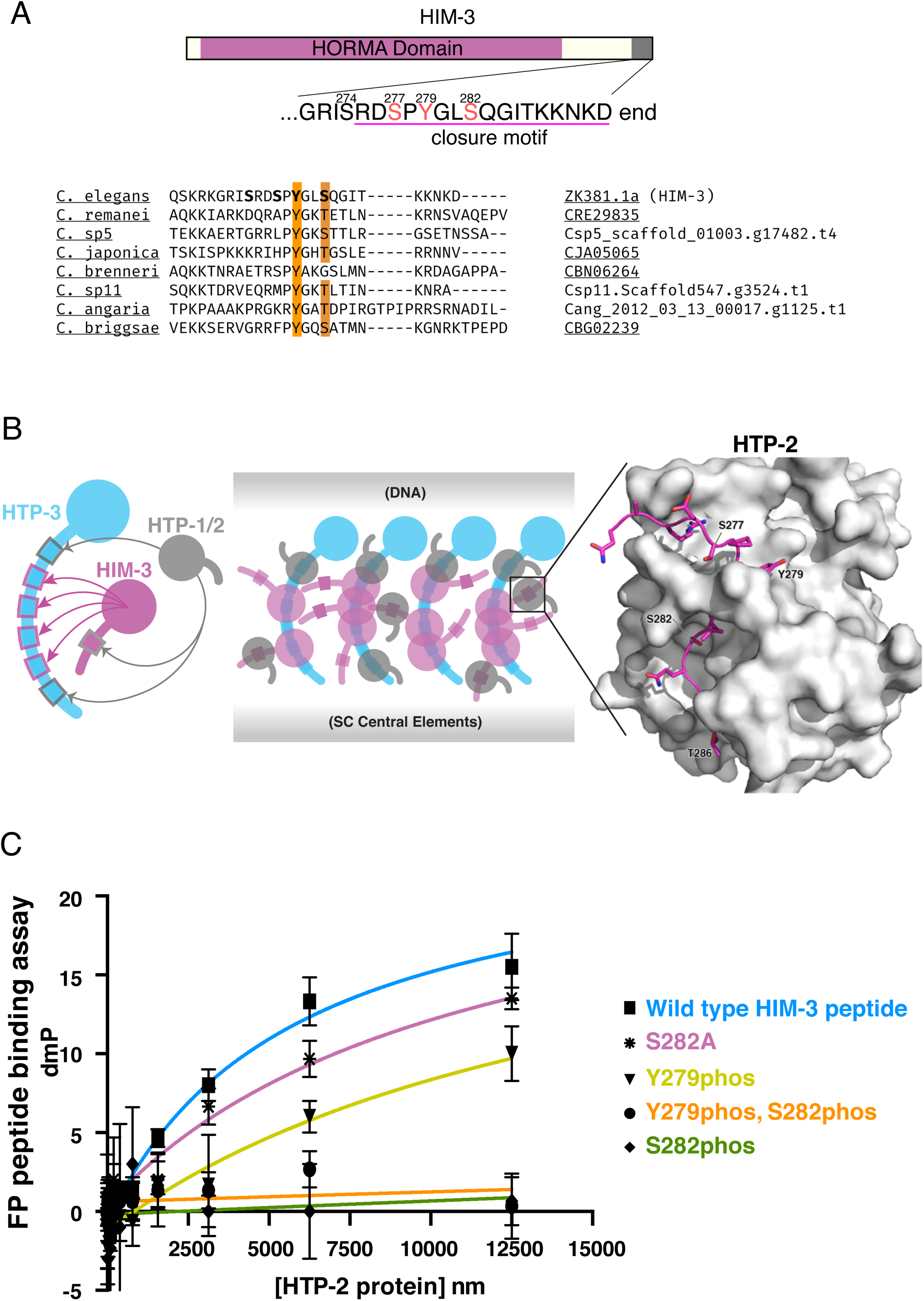
Phosphorylation at the HIM-3 closure motif prevents HTP-1/2 binding *in vitro*. **A**, *top:* Schematic diagram of HIM-3, showing the HORMA domain (magenta) and phosphorylation sites identified by mass spectrometry in red. The conserved closure motif is underlined. *bottom:* Conservation of the Tyr and Ser/Thr, highlighted in orange, in the closure motif in eight *Caenorhabditis* species. **B**, *left:* diagram of hierarchical binding of HTP-1, 2, 3 and HIM-3 from [24]. *center*, diagram of possible multivalent binding modes of the axial element, based on [33,66]. *right*, Interactions between the HIM-3 closure motif (magenta) and the HTP-2 (grey) HORMA domain shown in PDB structure 4TZL [24]. **C**, Fluorescence polarization peptide binding assay using bacterially purified HTP-2 protein (full length) and fluorescently tagged HIM-3 peptides with indicated phosphorylation or A1a substitution. Wild type means HIM-3 peptides with no phosphorylation. Phosphorylation at Ser282 inhibits HTP-2 binding *in vitro*. Error bars are calculated as the SD from duplicated measurements. Measured Kd values were: 5.8µM for unphosphorylated peptides, not significant binding for S282phos and Y279phos_S282phos peptides, 16.1µM for Y279phos peptides, and 11.3µM for S282A peptides.

### Phosphorylated HIM-3 localizes to the entire length of the SC and then becomes enriched on the short arms

To further understand the function of HIM-3 phosphorylation, we generated phospho-specific antibodies against HIM-3 peptides containing phosphorylated Ser277 and Ser282 since we detected this doubly-phosphorylated peptide most frequently by mass spectrometry. Lack of staining with this antibody in *him-3* mutants in which both Ser277and Ser282 are converted to A1a confirmed specificity for the phosphoresidues (**Supplemental Figure 1B**). Immunofluorescence using HIM-3phos specific antibodies showed that phosphorylation of HIM-3 starts from the beginning of meiotic prophase. From leptotene through mid-pachytene, HIM-3phos antibody staining coincides with that of “panHIM-3” antibodies that were raised against the entire protein and are thus expected to recognize both phosphorylated and unphosphorylated HIM-3 (**Figure 2A**). As oocyte precursor cells progress into late pachytene, designate crossovers, and begin to differentiate long and short arms, HIM-3phos antibody staining becomes enriched on only a small, terminal part of the SC, in contrast to panHIM-3 staining which remains on the full length of the SC, suggesting that phosphorylated HIM-3 becomes enriched on the short arm (**Figure 2B**). Simultaneous staining with the short arm marker SYP-1phos antibody [16] as well as the crossover designation marker COSA-1 [32] revealed that phosphorylated HIM-3 becomes enriched at short arms once crossovers are designated (**Figure 2C**). Although Ser282 is in an ATM/ATR kinase consensus site (SQ), we found that phosphorylation of HIM-3 Ser282 does not depend on these DNA damage-responsive kinases (**Supplemental Figure 1C**).

**Figure 2.**
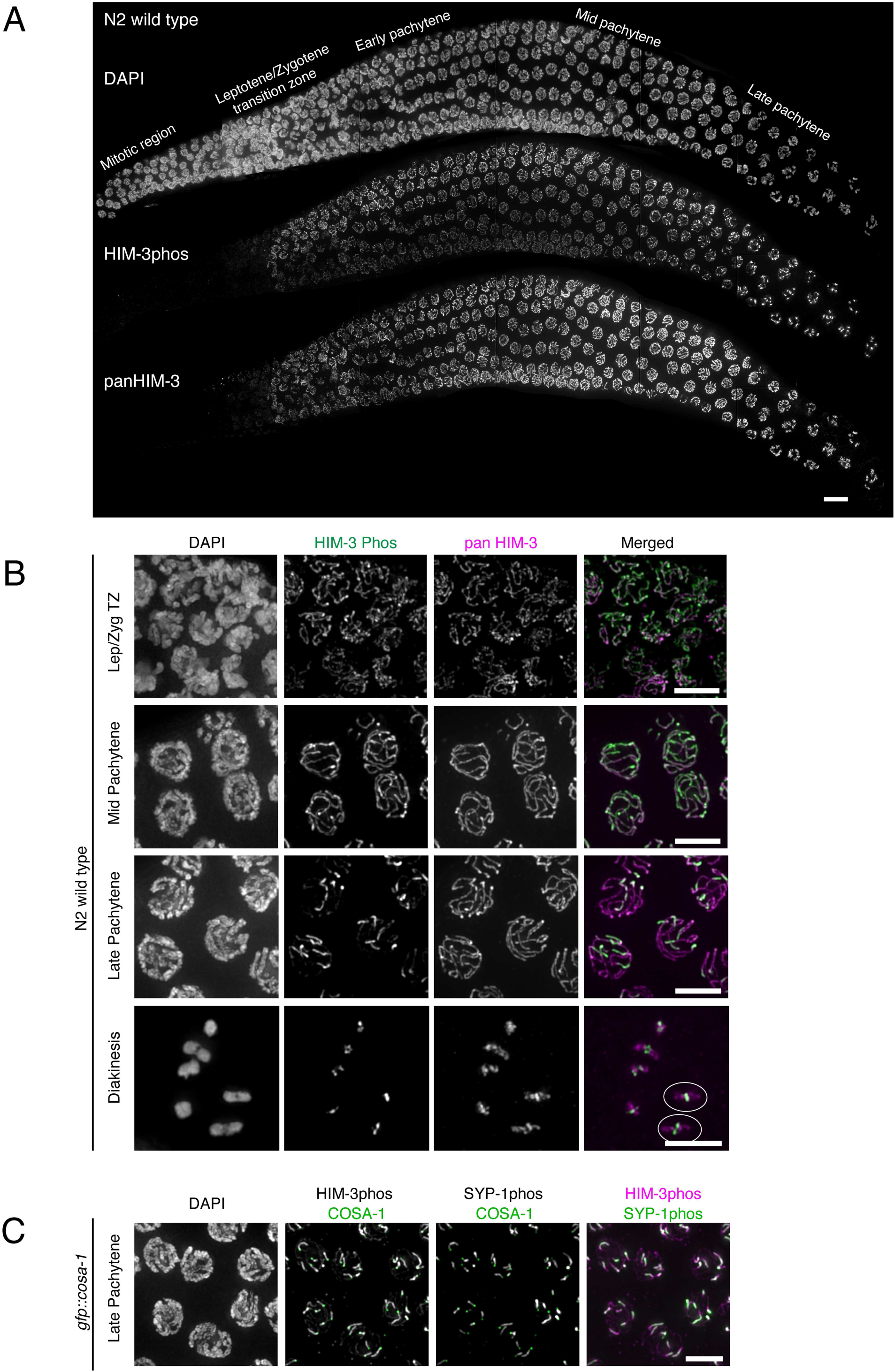
Phosphorylated HIM-3 localizes to the SC and becomes enriched on short arms. **A**, wild-type germline showing DNA stained with DAPI (*top*), and immunostaining with phosphospecific HIM-3 antibodies (*middle*), and non-phosphospecific HIM-3 (pan-HIM-3) antibodies (*bottom*). **B**, Wild-type N2 oocyte precursor cells taken at different meiotic stages and immunostained for phospho-HIM-3 (green in merged images) and pan-HIM-3 (magenta in merged images). Diakinesis chromosomes with cruciform HIM-3 staining are circled in white. **C**, Late pachytene cells (*gfp::cosa-1*) co-immunostained for phospho-HIM-3 (magenta in merged image), phospho-SYP-1 (green in merged image), and anti-GFP showing the position of GFP-COSA-1 (green in two middle images). Scalebars, 5µm.

Compared to the complete confinement of phosphorylated SYP-1 to short arms immediately after meiocytes enter late pachytene, phosphorylated HIM-3 enrichment was less complete and began slightly later (**Figure 3A**). nrichment of phosphorylated HIM-3 was detected at the very end of late pachytene, and weak HIM-3phos staining was still detectable on long arms until phosphorylated HIM-3 became completely confined to short arms at −1 diakinesis, the most mature oocyte precursor cell (proceeding distally from the spermatheca, oocyte precursors in diakinesis are designated as stage −1, −2, −3 etc. oocytes)(**Figure 2B**). In addition to restriction of localization, HIM-3phos staining appeared to increase in intensity once partitioning was observed. To quantitatively measure this, we compared the levels of phosphorylated HIM-3 on short arms after partitioning with that of the entire chromosome axis before partitioning within the same gonad. Dividing late pachytene nuclei into pre- and post-partitioned regions, we picked 500 points on the SC in each region and plotted the intensity value per pixel after normalizing to the average intensity values of the pre-partitioning region. Our quantification showed that HIM-3phos signal intensity per pixel is higher on short arms after partitioning than along the entire axis before partitioning, indicating a net increase of phosphorylated HIM-3 on short arms (approximately 1.5 times increase, n=6 gonads)(**Figure 3B, Supplemental Figure 2A, B, C**). In addition, we compared panHIM-3 staining levels before and after HIM-3phos partitioning as well as short versus long arms after partitioning. We also found a slight (approximately 1.2 times increase, n=6 gonads) but significant increase of panHIM-3 intensity on short arms after partitioning, suggesting a net addition of HIM-3 proteins to the SC toward the end of pachytene (**Figure 3B**). This is consistent with the previous observation that HIM-3 and HTP-1/2 continue to accumulate on the SC throughout pachytene and diplotene [33]. In nuclei with partitioned HIM-3phos, panHIM-3 staining levels did not differ between short and long arms, suggesting that the chromosome-wide increase in panHIM-3 levels does not derive from preferential addition of HIM-3 proteins to short arms but rather global addition along the entire SC (**Figure 3C**). We reasoned that these observations could be interpreted in one of the following ways: (1) global dephosphorylation of HIM-3 with counter-acting phosphorylation specifically on short arms or (2) long arm-specific dephosphorylation and short arm-specific phosphorylation of HIM-3 (**Figure 3D**). Although other scenarios such as removal of phosphorylated HIM-3 proteins specifically from long arms with additional phosphorylation on short arms, or dephosphorylation from long arms and addition of phosphorylated HIM-3 on short arms are conceivable, we disfavor these possibilities since we did not detect any change in pan-HIM-3 staining levels between short and long arms after partitioning. However, we cannot exclude the possibility that the fraction of phosphorylated HIM-3 may be relatively small compared to the entire pool of axis-bound HIM-3, making the removal or addition of phosphorylated HIM-3 proteins undetectable by pan-HIM-3 staining. Since a previous photoconversion experiment showed that HIM-3 is rather static and stable once incorporated to the chromosome axis [34], we judge it unlikely that phosphorylated HIM-3 moves from long to short arms. Altogether, our data suggest that short arm enrichment of HIM-3 is likely to occur through a combination of dephosphorylation of phosphorylated HIM-3 from long arms and short arm-specific phosphorylation activity.

**Figure 3.**
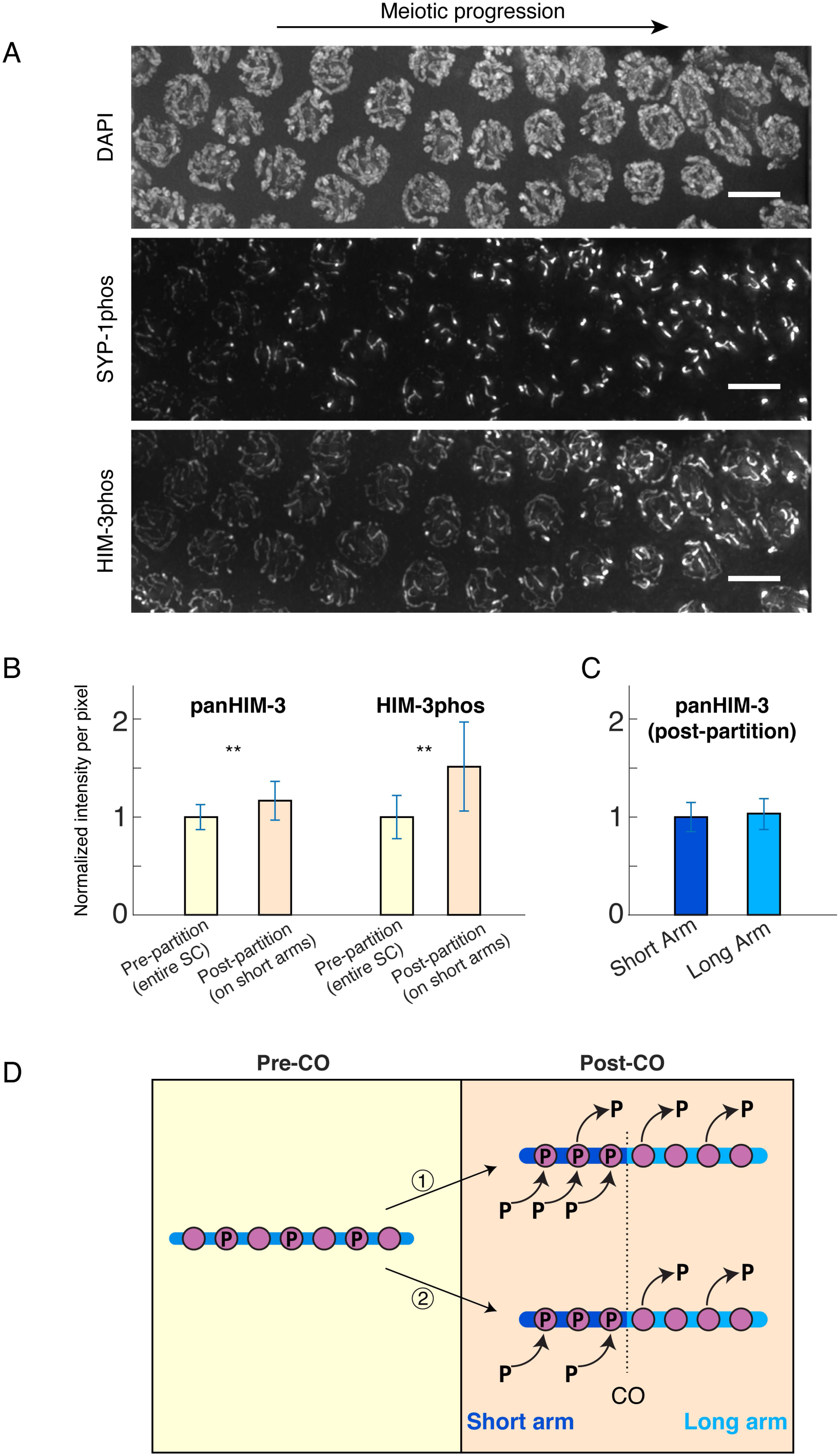
HIM-3 phosphorylation increases on the short arm, while pan-HIM-3 increases slightly along the whole chromosome. **A**, a segment of a wild-type germline (mid to late pachytene) showing meiotic progression from left to right. The accumulation of SYP-1 phosphorylation to one region (the short arm) can be seen to precede and exceed the partitioning of HIM-3. Scalebars, 5µm. **B**, quantitation of pan-HIM-3 and phospho-HIM-3 staining in pre-partitioned chromosomes (yellow bars), post-partitioned chromosomes (beige, right). For both epitopes, values for post-partitioned chromosomes are significantly greater than that for pre-partitioned chromosomes (two-tailed T test, p<0.00l). 3000 points for each condition (pre and post) are compared (50 points in each of l0 nuclei from 6 gonads); full point distributions for each set are shown in **Supplemental Figure 2A, B and C**. **C**, normalized intensity values of pan-HIM-3 in late pachytene nuclei comparing the short with the long arm. No significant difference exists between short and long arm staining (two-tailed T test). 3000 points for each condition (short and long) are compared (50 points in each of 10 nuclei from 6 gonads). Short arm points are the same points used for “post-partitioning” in **B**. For both **B** and **C**, intensity values derive from dividing raw pixel intensities of all individual pre- and post-partitioned points by the mean intensity of pre-partitioned points in the same gonad. Bars show mean values and error bars show standard deviation. **D**, illustration of the two scenarios for phospho-HIM-3 enrichment discussed in the text. The blue line is the chromosome; violet circles stand for HIM-3 proteins; “P” indicates phosphorylation; arrows indicate the adding or leaving of molecules. On the right side, the short arm (dark blue) is toward the left of the vertical line indicating the crossover position (marked with “CO”); the long arm (light blue) is toward the right.

Since the activity of PP1 phosphatase, GSP-2, has been shown to localize to long arms upon crossover designation via HTP-1/2 and LAB-1 interactions [18,21], we next tested whether GSP-2^PP1^ dephosphorylates HIM-3 on long arms by immunofluorescence. In *gsp-2* mutants, HIM-3-phos was still enriched on short arms and absent from long arms, showing no difference from wild-type staining, suggesting that GSP-2^PP1^ is not responsible for loss of phosphorylated HIM-3 from long arms (**Supplemental Figure 3A**).

To understand the mechanism of the putative kinase activity on short arms, we examined the localization of phosphorylated HIM-3 in various meiotic mutant backgrounds. Unexpectedly, we found that HIM-3 phosphorylation is dependent on synapsis (**Supplemental Figure 3B**). In *syp-1(me17)* null mutants or other meiotic mutants such as *plk-2(ok1936)* that generate unsynapsed chromosomes [35-37], HIM-3phos staining was absent from unsynapsed chromosomes. This suggests that recruitment or activation of one or more kinases that phosphorylate HIM-3 is dependent on SYP proteins. The enrichment of SC central proteins on short arms in late pachytene [4,38-41], coincident in location and timing with phosphorylated HIM-3 partitioning, further suggests the central element provides a platform for a kinase to phosphorylate HIM-3 preferentially on short arms upon crossover designation.

### Asymmetric enrichment of phosphorylated HIM-3 is dependent on crossover designation

Next, we tested if asymmetric enrichment of phosphorylated HIM-3 is dependent on the formation of crossover intermediates. DNA double strand breaks (DSBs) are created by SPO-11 a topoisomerase-like enzyme, to initiate homologous recombination. In *spo-11(me44)* mutants, no programmed DSBs are generated during meiotic prophase and thus meiocytes fail to form crossovers [42]. In *spo-11 (me44)* mutants, the great majority of nuclei lack both DSBs and crossover designation marker COSA-1, and show HIM-3phos staining along the entire length of the SC even at very late pachytene **(Figure 4A)**. To further test whether crossover intermediates *per se* and not only DSBs are required for enrichment of phosphorylated HIM-3, we visualized phospho-HIM-3 and SYP-2 in *msh-5* mutants, which achieve even higher DSB levels than wild-type, but do not form crossover intermediates [43,44]. We found no partitioning of phospho-HIM-3 in *msh-5* mutants, indicating that crossover intermediates containing MSH-5 are required for the enrichment (**Figure 4C**).

**Figure 4.**
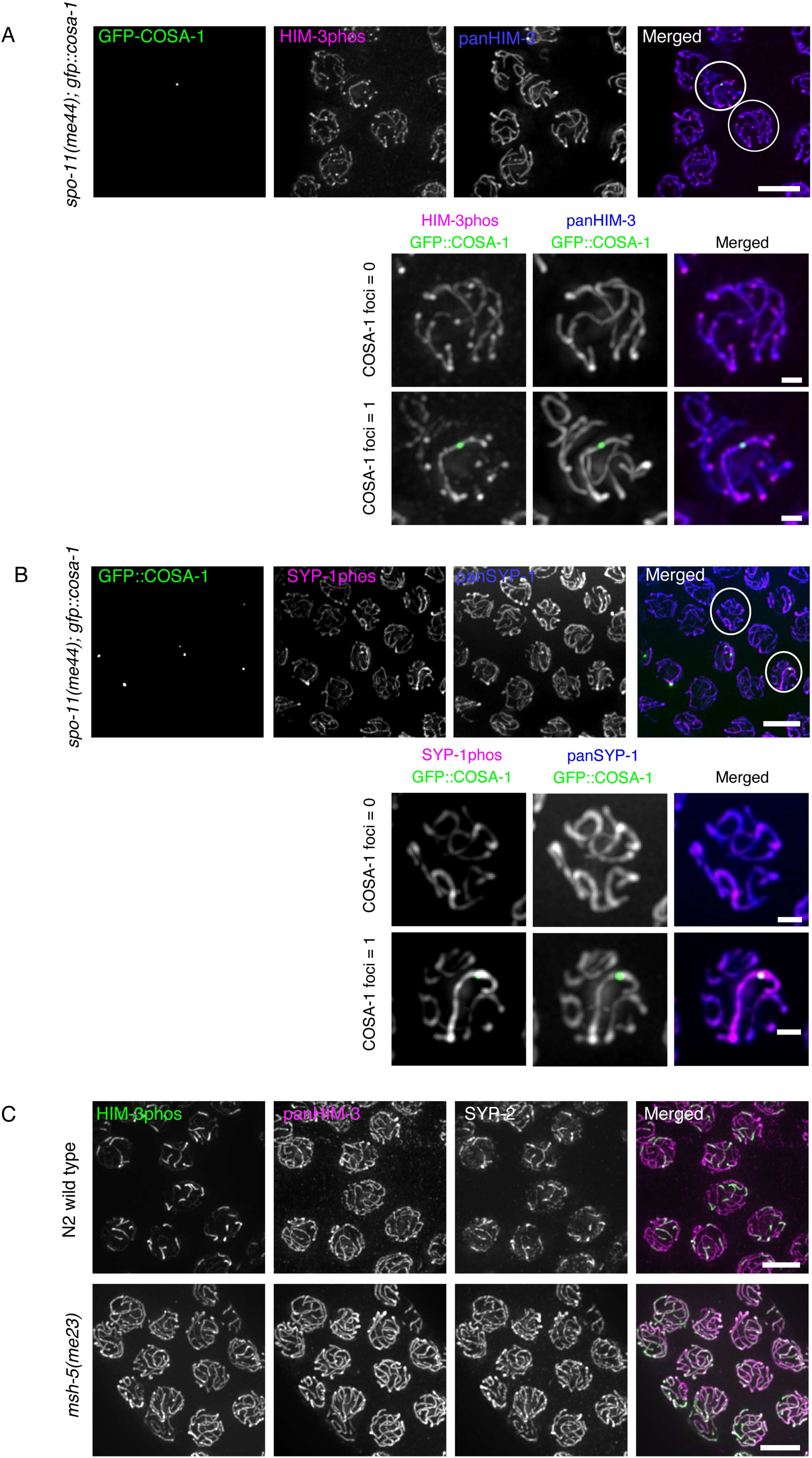
Asymmetric localization of phosphorylated HIM-3 requires crossover intermediates. **A**, Immunostaining of crossover designation marker GFP-COSA1 (green in merged image), phospho-HIM-3 (magenta in merged image), and pan-HIM-3 (blue in merged image) in *spo-11(me44); gfp-cosa-1* oocyte precursors. **B**, Immunostaining of GFP-COSA1 (green in merged image), phospho-SYP-1 (magenta in merged image), and pan-SYP-1 (blue in merged image) in *spo-11(me44); gfp-cosa-1* oocyte precursors. For both **A** and **B**, at top right, white circles surround nuclei shown magnified in insets below; COSA-1 focus number (0 or 1) is indicated. **C**, wild-type (top row) and *msh-5* (bottom row) germlines at late pachytene stained with antibodies against phospho-HIM-3 (green in merged images), pan-HIM-3 (magenta in merged images), and SYP-2 (gray; not shown in merged images). Scalebars: 5µm (unmagnified), lµm (magnified).

Previous studies showed that a small minority of chromosomes in *spo-11 (me44)* mutants contain COSA-1 foci, which presumably result from unrepaired DNA damage incurred during replication; however, these foci are not able to mature into genuine crossovers [45]. The same study also has shown that chromosomes with such COSA-1 foci become strongly enriched for SYP-1 and its interacting protein PLK-2 in a chromosome-autonomous manner, at the expense of chromosomes that lack COSA-1 foci [45]. Similarly, we found that rare chromosomes with COSA-1 foci in *spo-11(me44)* mutants show enrichment of phosphorylated HIM-3 along the entire length of the chromosome. In these same nuclei, HIM-3phos staining was mostly absent from chromosomes lacking COSA-1. However, pan-HIM-3 staining intensity appeared the same between chromosomes with and without COSA-1 foci. Taken together, these observations suggest that HIM-3 is phosphorylated on chromosomes bearing COSA-1 foci, and dephosphorylated on chromosomes without COSA-1.

Next, we checked whether phosphorylation of other SC components displayed a similar pattern to HIM-3phos in nuclei with a single COSA-1 focus. We found that phosphorylated SYP-1 at Thr452 also becomes enriched along the entire length of chromosomes harboring COSA-1 foci in *spo-11(me44)* before the bulk of SYP-1, which was visualized by pan-SYP-1 antibody (**Figure 4B**). This is consistent with a previous observation that PLK-2 accumulates on chromosomes with DNA breaks, and with our previous evidence suggesting that phosphorylated SYP-1 at Thr452 recruits PLK-2 to the SC [16,45]. These results suggest that the presence of recombination intermediates containing MSH-5 (i.e., crossover intermediates in wild-type animals) alters the chromosome environment in *cis*, and enrichment of phosphorylated HIM-3 and SYP-1, compared to their unphosphorylated forms, is more responsive to this chromosome-wide change.

### Asymmetric enrichment of phosphorylated HIM-3 is sensitive to the number of crossovers

The observed enrichment of phosphorylated HIM-3 and SYP-1 in rare *spo-11* nuclei that contain a single chromosome with a COSA-1 focus differs from the wild-type situation in one important respect: both phosphoproteins fail to partition to the short arm of the chromosome, but rather remain present along the entire chromosome length. We reasoned this difference could be due either to the differing physical nature of DSBs induced by unrepaired mitotic damage in *spo-11* mutants, compared to SPO-11-catalyzed DSBs, or to the lower number of crossover intermediates present. To distinguish these possibilities, we examined *dsb-2* mutants, in which DSBs are still catalyzed by SPO-11 but are present at greatly reduced levels, giving rise to meiocytes with varying numbers of DSBs and thus crossovers (ranging from 0 to 6 COSA-1 foci) in the same gonad [46]. In *dsb-2* mutants, we observed partitioning of HIM-3phos to short arms, but only in nuclei with relatively high numbers of crossover intermediates. Complete partitioning of HIM-3phos was detected only in nuclei with 5 or 6 COSA-1 foci in late pachytene, whereas accumulation along the entire length of chromosomes was seen in nuclei with 4 or fewer COSA-1 foci. The fraction of nuclei with partial (some but not all chromosomes with a COSA-1 focus achieved partitioning) or complete partitioning of HIM-3phos increased as meiotic prophase progressed, and in diplotene, about 20% of nuclei with 4 COSA-1 foci also achieved complete partitioning (**Figure 5A,C**). In contrast, complete partitioning of SYP-1phos was already achieved in late pachytene in many nuclei with 4 or more COSA-1 foci (44% of nuclei with CO=4, and 73% of nuclei with CO=5), and no partitioning was observed in nuclei with 1, 2 or 3 COSA-1 foci (**Figure 5B,C**). Since SYP-1 disassembles completely from long arms in diplotene, our quantification of SYP-1phos partitioning was limited to late pachytene nuclei but did not include diplotene. While we frequently observed a high proportion of partial partitioning of HIM-3phos, partial partitioning of SYP-1phos was relatively rare, suggesting that partitioning of SYP-1phos is a more rapid, switch-like process involving feedback mechanisms. These results show that the overall number of crossover intermediates in a nucleus can cumulatively influence the efficiency of short-versus-long arm partitioning.

**Figure 5.**
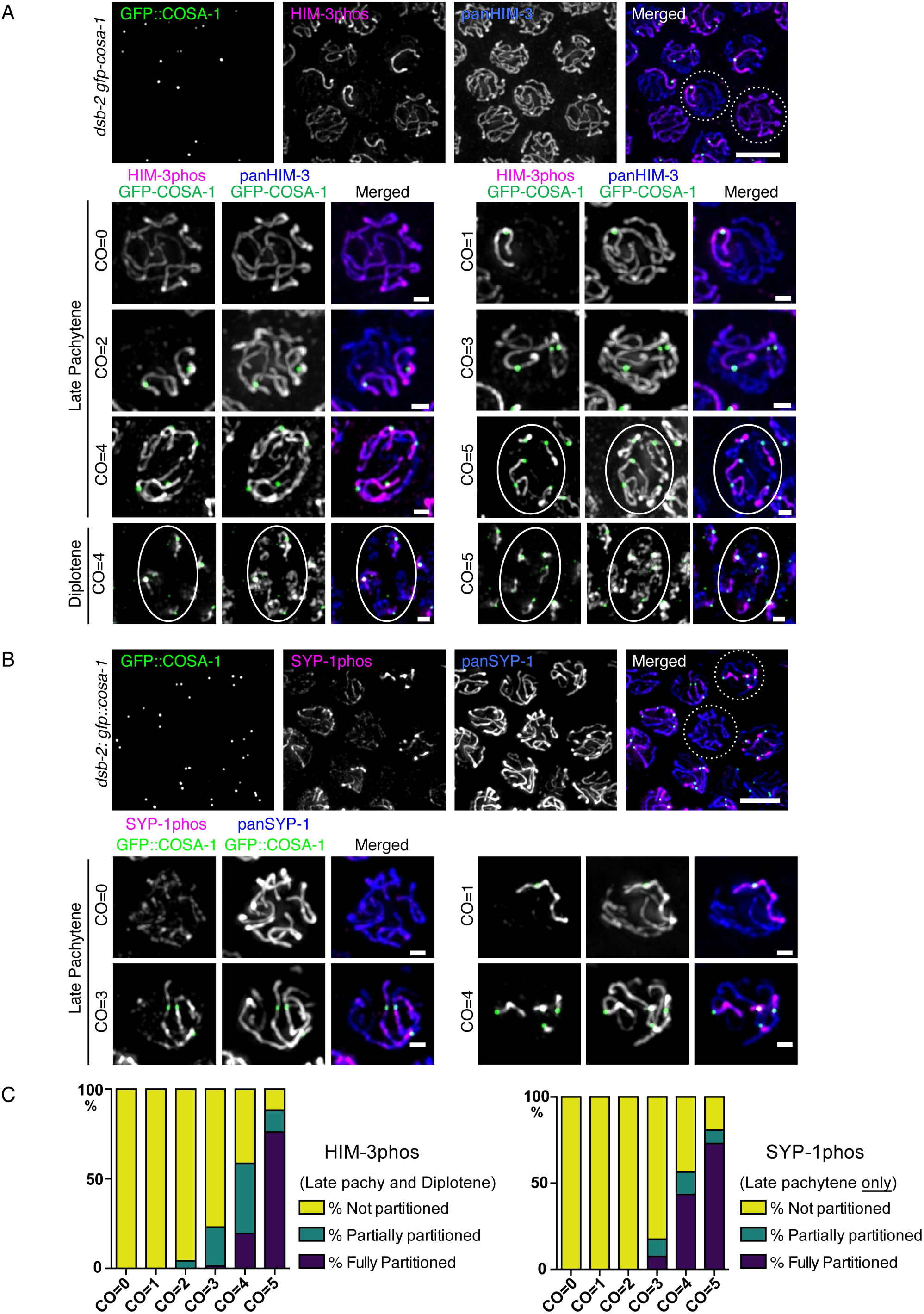
Asymmetric localization of phosphorylated HIM-3 depends on the number of crossover intermediates. **A**, *Top:* Immunostaining of crossover designation marker GFP-COSA1 (green in merged image), phospho-HIM-3 (magenta in merged image), and pan-HIM-3 (blue in merged image) in *dsb-2(me96); gfp-cosa-1* oocyte precursors (late pachytene). Dotted-circled nuclei are shown magnified in the top row below. *Bottom*, individual nuclei with different numbers of COSA-1 foci (left labels). Ellipses surround those nuclei with asymmetrically partitioned phospho-HIM-3. **B**, *Top:* Immunostaining of GFP-COSA1 (green in merged image), phospho-SYP-1 (magenta in merged image), and pan-SYP-1 (blue in merged image) in *dsb-2(me96); gfp-cosa-1* oocyte precursors (late pachytene). *Bottom*, individual nuclei with different numbers of COSA-1 foci (left labels). Scalebars: 5µm (overview images), 1µm (individual nuclei). **C**, quantitation of observed nuclei stained with either phospho-HIM-3 (left, n=243) or phospho-SYP-1 (right, n=l64) showing classification of partitioning.

### Phosphoregulation of HIM-3 promotes timely SC disassembly

To understand the function of HIM-3 phosphorylation, we used CRISPR to generate non-phosphorylatable as well as phospho-mimetic *him-3* mutants at the original locus. For non-phosphorylatable mutants, Ser277, Tyr279 and Ser282 were respectively mutated to A1a, Phe and A1a either solely (S282A mutant) or in combination (FA mutant: Y279F+S282A; AFA mutant: S277A+Y279F+S282A). In a fluorescence polarization assay, HIM-3 peptides carrying the S282A point mutation retained HTP-2 binding capacity close to wild type (**Figure 1C**). For phosphomimetic mutants, Ser282 was mutated either to Glu or Asp (S282 or S282D mutant). Some of the point mutations were generated in the background of C-terminally FLAG-tagged *htp-1* (*htp-1::flag)* to facilitate HTP-1 visualization by immunofluorescence. Adding a FLAG tag did not confer any meiotic defects as this strain showed normal embryonic viability (**Supplemental Figure 4A**). In all of the *him-3* mutants, HIM-3 proteins localize to the SC at normal levels by pan-HIM-3 antibody staining. In *him-3* phosphomimetic mutants, HTP-1 was detected on the SC from the leptotene/zygotene transition one, and SC central elements were polymerized normally. However, as expected for phosphomimetic mutants,quantitative image analysis showed that global HTP-1::flag levels in *him-3(S282E) htp-1::flag* are reduced in diakinesis compared to control gonads *(htp-1::flag* by itself) dissected and immunostained on the same slide (**Supplemental Figure 4B**). This implies that although HTP-1 may bind to the closure motif of either HIM-3 or HTP-3 [24], some fraction of HTP-1 does normally bind to HIM-3, presumably the non-phosphorylated pool, *in vivo*. Since the fluorescence polarization assay suggested HIM-3 S282 phosphorylation prevents HTP-1/2 binding, and phosphorylated HIM-3 is enriched on short arms from which HTP-1/2 is removed, we wondered whether HIM-3 phosphorylation prevents HTP-1/2 from re-binding to short arms once they dissociate from the SC. To test this, we examined the confinement of HTP-1 to long arms as well as confinement of SYP-1 to short arms in diakinesis nuclei in *him-3* phospho-mutants. Since condensed chromosomes at diakinesis are difficult to resolve with light microscopy due to the arbitrary orientation of chromosome arms relative to the optical axis, we limited our analysis to bivalent chromosomes in which we could clearly resolve a cruciform structure of HTP-3, which localizes to both short and long arms. Within each nucleus, we scored as many resolvable bivalent chromosomes as possible, and counted the nucleus as positive if at least one bivalent showed abnormal localization of SYP-1 or HTP-1/2. Since SYP-1 dissociates from chromosomes and cannot be detected reliably in −3, −2 and −1 diakinesis nuclei in wild type, the analysis was limited to −7 through −4 diakinesis nuclei. If HIM-3 phosphorylation normally functions to prevent HTP-1/2 from re-binding to short arms, non-phosphorylatable *him-3* mutants would be expected to retain HTP-1/2 on short arms. However, we did not detect any delay in HTP-1 dissociation from short arms in *him-3(S282A)* or *(AFA)* mutants (**Figure 6A,B,C**, only quantification is shown for S282A). To our surprise, instead, phosphomimetic mutants (S282D or S282) delayed HTP-1 dissociation from short arms in about 25% of oocytes, which show persistent HTP-1 on short arms in early diakinesis, while HTP-1 became completely restricted to long arms during the earlier diplotene stage in control gonads (**Figure 6**). Although HTP-1 persisted longer on short arms in *him-3* phosphomimetic mutants, eventually HTP-1 was confined to long arms in −1 diakinesis nuclei, suggesting that partitioning of phosphorylated HIM-3 influences the timing of HTP-1 dissociation from short arms but is not absolutely required. Unexpectedly, both non-phosphorylatable and phosphomimetic *him-3* mutants also delayed SYP-1 confinement to short arms and showed persistent SYP-1 on long arms in the majority of nuclei through diakinesis, while in the wild type SYP-1 is completely confined to short arms in early diplotene. Although SYP-1 persisted on both arms in many *him-3* mutant nuclei, SYP-1 eventually disappeared from chromosomes in late diakinesis nuclei with normal timing (−3 diakinesis and onward). Our results suggest that partitioning of phosphorylated HIM-3 to short arms promotes global rearrangement of SC components, including central elements, along the entire length of the SC rather than locally on short arms. Further, the normal dissociation of HTP-1 from the short arm in *him-3* non-phosphorylatable mutants shows that HIM-3 phosphorylation is not required to prevent HTP-1/2 from re-binding the short arm in diplotene after their dissociation.

**Figure 6.**
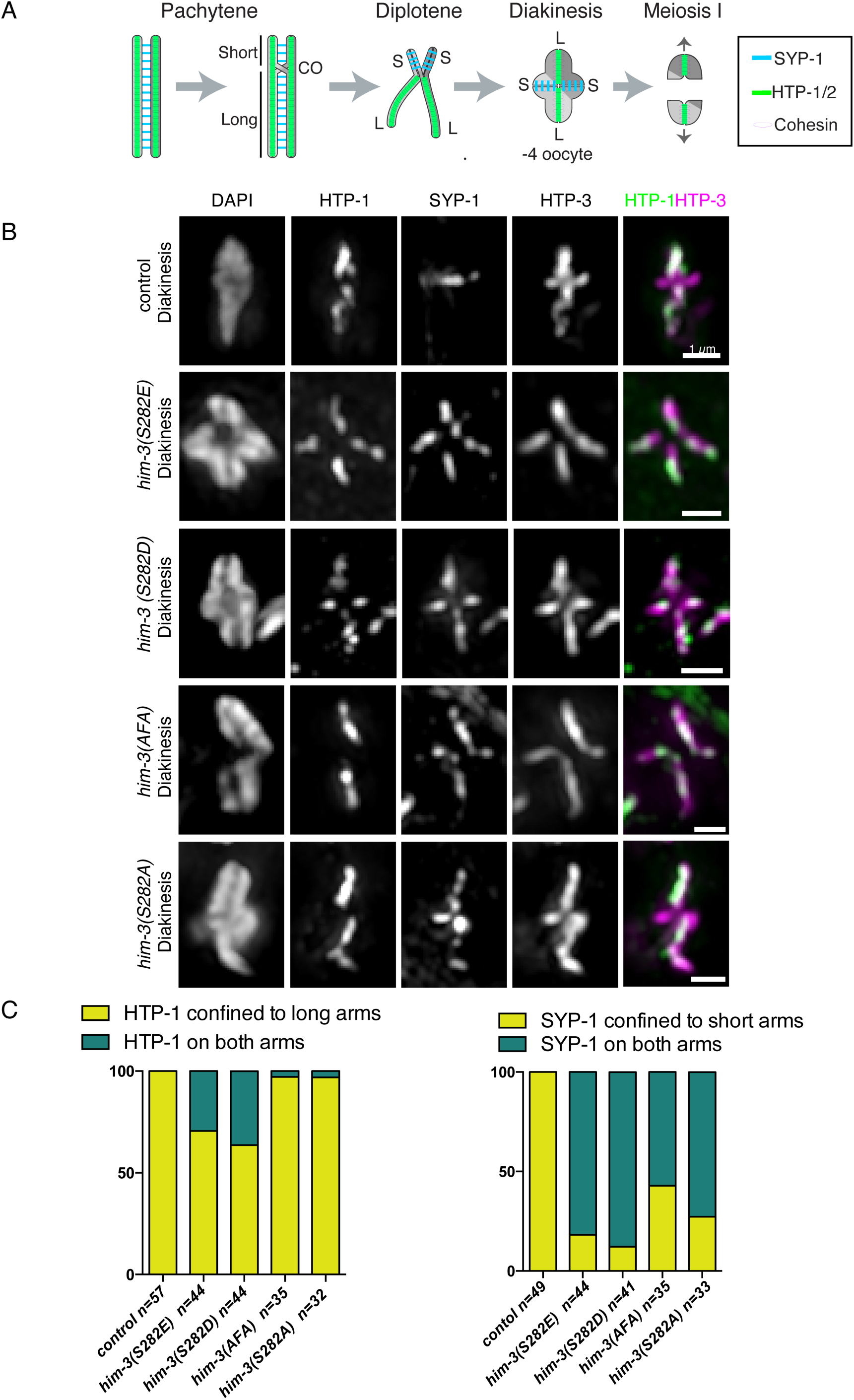
Phosphoregulation of HIM-3 is involved in timely SC disassembly. **A**, diagram of chromosome remodeling after crossover designation, showing normal localization of SYP-1 and HTP-1/2 at diakinesis. **B**, Immunostaining of HTP-1 (green in merged images), SYP-1, and HTP-3 (magenta in merged images) on −4 to −7 diakinesis nuclei of *htp-1::flag, him-3(S282E) htp-1::flag, him-3(S282D), him-3(AFA) htp-1::flag*, and *him-3(S282A) htp-1::flag*. Single chromosome images were cropped out of 3D data sets and shown as partial projections. Scalebars, 1µm. **C**, quantitation of partitioning degree of HTP-1 (left) and SYP-1 (right) for *htp-1::flag, him-3(S282E) htp-1::flag, him-3(S282D), him-3(AFA) htp-1::flag and him-3(S282A) htp-1::flag*. The number of diakinesis nuclei scored is shown beneath the corresponding bar.

Since asymmetric SC disassembly is a precursor to establishment of chromosome separation domains, and previous studies on mutations that perturb asymmetric SC disassembly also mislocalized CPC activity [4,15,16,18,29], we examined CPC localization on short arms by immunofluorescence of CPC component ICP-1 (the *C. elegans* INC NP ortholog) in *him-3* mutants. In wild type, ICP-1 is detected in −3, −2, −1 diakinesis nuclei and later, is strongly enriched on short arms in −3 and −2 diakinesis, and becomes fully confined to short arms from −1 diakinesis through metaphase I oocyte pronuclei (**Supplemental Figure 5**). In *him-3(S282D)* mutants, ICP-1 short arm confinement was reduced, but S282A, AFA or S282 mutants did not show a significant difference from control animals. Also, consistent with ICP-1 localization, embryonic viability and the percentage of male progeny (indicative of *X* chromosome meiotic nondisjunction) are very similar to wild type in non-phosphorylatable mutants, and only a slight increase in embryonic lethality and male progeny number are observed in phospho-mimetic mutants (0.8% embryonic lethality in control v.s. 5.1% lethality in S282; 0.3% lethality in control v.s. 2.2% lethality in S282D: **Supplemental Figure 4A**). This suggests that although SYP-1 dissociation is significantly delayed in all *him-3* phospho-mutants, and asymmetric distribution of HTP-1 is delayed in *him-3* phospho-mimetic mutants, this does not lead to gross perturbation in designating chromosome separation sites, perhaps because other pathways including ZHP-1/2, phosphorylated SYP-1 and PLK-2 [15], akirin [47], or MAP kinase and phosphorylated SYP-2 [17] redundantly function to designate the separation site.

### Partitioning of phosphorylated HIM-3 depends on SYP-1 Thr4S2 phosphorylation

Since the timing of phosphorylated HIM-3 enrichment on short arms follows confinement of phosphorylated SYP-1, we next asked whether HIM-3phos enrichment is a downstream event of SYP-1phos confinement or a parallel pathway for asymmetric SC disassembly. In non-phosphorylatable *syp-1(T452A)* mutants or *plk-2(ok1936)* mutants, HIM-3 was phosphorylated normally on synapsed chromosome regions beginning at the leptotene/zygotene transition one, but phosphorylated HIM-3 failed to accumulate on short arms in late pachytene (**Supplemental Figure 6A**). This suggests that while initial phosphorylation of HIM-3 does not require SYP-1 phosphorylation at Thr452 and subsequent PLK-2 loading, partitioning of phosphorylated HIM-3 to the short arm does require SYP-1 phosphorylation. In contrast, in *him-3(S282E)* or (*AFA*) mutants, phosphorylated SYP-1 was partitioned normally to short arms (**Supplemental Figure 6B**). These observations suggest that partitioning of phosphorylated HIM-3 is a downstream event of phosphorylated SYP-1 partitioning, and in turn, promotes dissociation of SYP-1 from long arms. This observation shows that partitioning of SYP-1 phosphorylation to the short arm is not sufficient to remove SC central element proteins from the long arm.

Somewhat unexpectedly, we noted that in −4 and −3 diakinesis nuclei, weak phosphorylation of SYP-1 was observed on long arms in *him-3* phos-phomutants, although it was strictly limited to short arms at earlier time points (**Supplemental Figure 6B**). Since SYP-1 normally begins to dissociate from the chromosome axis at −4 diakinesis, it is unlikely that phosphorylated SYP-1 proteins are newly added to long arms in these mutant nuclei. Rather, hyperpersistent SYP-1 on long arms in *him-3* phospho-mutants is likely phosphorylated *de novo* by a kinase that is activated only in mature diakinesis nuclei. In the wild type, since SYP-1 is already limited to short arms by diakinesis, this kinase activity would not lead to accumulation of phosphorylated SYP-1 on long arms. Altogether, our results suggest that HIM-3phos partitioning is a downstream event of SYP-1phos partitioning. Since disruption of SYP-1 phosphorylation leads to stronger perturbations in HTP-1/2 localization, CPC localiation and the first meiotic division [16] than seen in *him-3* phospho-mutants, we conclude HIM-3phos partitioning is one of many downstream events depending on SYP-1phos partitioning.

### Phosphoregulation of HIM-3 is not required to prevent recombination via sister chromatids nor to satisfy the synapsis checkpoint

During meiotic prophase, programmed DSBs are preferentially repaired via inter-homolog recombination to generate crossovers between homologs while inter-sister recombination is actively suppressed by a mechanism depending on, among other proteins, HIM-3 and HTP-1/2 [26,48]. In various organisms, HORMA domain proteins are required for homolog-biased recombination, and specifically phosphorylation of the budding yeast HORMA domain protein, Hop1, by ATM/ATR kinases has been shown to be important for inter-homolog bias [49-53]. We therefore wondered whether HIM-3 phosphorylation functions to prevent inter-sister recombination during meiotic prophase. In *C. elegans*, sister chromatid-mediated homologous recombination requires BRC-1 (the *C. elegans* ortholog of BRCA1) and its interactor BRD-1 (BARD1 ortholog) [54,55], and mutants defective in preventing inter-sister recombination during meiotic prophase synergistically increase embryonic inviability when combined with *brc-1* or *brd-1* mutants [56]. To test whether *him-3* phosphorylation might block the use of the sister chromatid as a repair template, we combined our non-phosphorylatable *him-3* mutation with *brc-1* and *brd-1* mutations by generating triple mutants, *brc-1(tm1145) brd-1(dw1); him-3 (S282A)* or *brc-1(tm1145) brd-1(dw1); him-3(AFA)*, and scored embryonic viability. However, combining these mutations did not result in synthetic embryonic lethality (**Supplemental Figure 7A**). Similarly, *him-3(AFA)* and *(S282E)* mutations had no effect on progeny viability of 75 Gy r-irradiated animals (**Supplemental Figure 7B**), showing it is not defective in sister chromatid-based DNA repair caused by excess DNA damage [56,57].

The dependence of HIM-3 phosphorylation on synapsis suggested the possibility of its acting to satisfy the synapsis checkpoint, which subjects cells that fail synapsis to programmed cell death [58] and depends on chromosome axis proteins, including HIM-3 and HTP-1/2 [58-60]. However, we saw no significant increase in apoptotic nuclei in *him-3 (S282A)* or *(Y279F S282A)* germlines compared to controls, indicating that this phosphorylation is not required to satisfy the synapsis checkpoint (**Supplemental Figure 7C**). Taken together, these results indicate that HIM-3 phosphorylation at the closure motif is not involved in either the regulation of inter-homolog bias for homologous recombination or in the synapsis checkpoint.

## Discussion

Here, we have identified and characterized phosphorylation sites within the closure motif of the conserved HORMA domain protein HIM-3. We showed that phosphorylated HIM-3 is enriched on short arms, and this partitioning contributes to the asymmetric disassembly of SYP-1 and HTP-1/2. Since *him-3* phospho-mutants show a delay in asymmetric SC disassembly, HIM-3 phosphorylation is not just a passive phosphorylation as a consequence of being unbound by HTP-1/2, but is likely a part of a redundant collection of mechanisms ensuring establishment of short and long arms prior to the stepwise cohesin degradation at meiotic divisions.

By preventing association of HTP-1/2 with HIM-3, phosphorylation of the HIM-3 closure motif may functionally modulate the axis structure. A previous study has shown that HTP-1/2 is able to bind either to HIM-3 or HTP-3’s closure motifs [24]. A more recent study has shown that on average two HTP-1/2 molecules and three HIM-3 molecules are present at every HTP-3 molecule, with substantial variation between HTP-3 molecules [33]. This indicates that closure motifs are rarely if ever occupied to their theoretical maximum; instead, many must remain unbound. The diverse, multivalent assembly that this scheme allows may be important to provide flexibility for regulating SC function.

In yeast and mice, SC disassembly upon pachytene exit is triggered by Polo kinase (Cdc5), DDK, and Aurora B kinase [61-64]. It is predicted that phosphorylation of SC components by these kinases leads to destruction of the SC by proteolysis, but the responsible phosphorylation sites on SC components and molecular mechanisms of SC disassembly have not been identified yet. A previous study has shown that disruption of binding between a human HORMAD protein, Mad2, and its binding partner CDC20 containing the closure motif requires unfolding of the N-terminal region of the HORMA domain by the AAA+ ATPase TRIP13, and thus requires energy consumption [65]. Currently, details of the molecular mechanism promoting dissociation of HTP-1/2 from HIM-3 or HTP-3 on short arms, or whether dissociation requires consumption of energy or proteolysis, are lacking. Since our *him-3* non-phosphorylatable mutants showed normal dissociation of HTP-1/2 from short arms, it is unlikely that HIM-3 phosphorylation at the closure motif functions to prevent re-binding of dissociated HTP-1/2 upon crossover designation. Rather, phosphomimetic mutations in HIM-3 delayed HTP-1 disassembly from short arms. If, as we speculate, HIM-3 phosphorylation prevents HTP-1/2 binding *in vivo*, HTP-1/2 will likely be incorporated to the axis solely by binding to HTP-3 closure motifs l and/or 6 in *him-3* phospho-mimetic mutants. In this scenario, we speculate that dissociation of HTP-1/2 may be less efficient when bound to HTP-3 than when bound to HIM-3, and HIM-3 may provide a binding site for HTP-1/2 to be readily disassembled upon crossover designation in wild type. A previous study has shown that disrupting the two HTP-1/2 binding sites of HTP-3 causes HTP-1/2 to bind exclusively to HIM-3 and leads to a delay in synapsis [24], suggesting the possibility that a fraction of HIM-3 protein must be unoccupied by HTP-1/2 for timely polymerization of SYP-1. HIM-3 phosphorylation at its closure motif may prevent saturation of HIM-3 by HTP-1/2 and thus may promote timely interactions between central and axial elements of the SC.

In addition to delayed HTP-1/2 disassembly, dissociation of unphosphorylated SYP-1 from the long arm is delayed in both *him-3* non-phosphorylatable and phospho-mimetic mutants. Although there has not been any direct evidence that central elements directly bind to HIM-3, previous studies suggest that HIM-3 is likely the interface between central elements and the chromosome axis: superresolution microscopy analysis has shown that among the chromosome axis proteins, HIM-3 localizes most closely to central elements [66]. Point mutations at the HIM-3 closure motif may affect interactions between HIM-3 and central elements, leading to prolonged synapsis in *him-3* phosphomimetic or non-phosphorylatable mutants.

Although SC disassembly is delayed in *him-3* phospho-mutants, this does not lead to gross perturbations of two-step cohesion loss and meiosis segregation, as only a slight reduction in embryonic viability was observed in *him-3*phosphomimetic mutants. This is consistent with previous results that a *him-3* C-terminal deletion lacking the entire closure motif does not show major meiotic defects [24]. This observation adds to the growing evidence that redundant mechanisms ensure CPC localization and limit cleavage of cohesin to short arms at meiosis I in the absence of asymmetric disassembly of the SC. We have previously shown that loss of SYP-1 Thr452 phosphorylation abrogates asymmetric dissociation of HTP-1/2 and SYP-1, yet reduces embryonic viability only mildly (39.7% reduction in *syp-1(T452A)*). Similarly, *syp-2* phosphomimetic mutants, in which approximately 50% of bivalents have persistent SYP-1 on both short and long arms in diakinesis, have only a l4.3% reduction in embryonic viability [17]. In these mutants, slight enrichment of SYP-1 on short arms and HTP-1/2 on long arms relative to the other arm may be sufficient to limit the CPC activity to short arms, since CPC localization is known to be regulated through both positive and negative feedback [67]. Alternatively, other mechanisms independent of asymmetric SC disassembly may exist to ensure concentration of CPC activity to short arms at meiosis I. For example, since metaphase I bivalents orient with the long arms pointing toward opposite spindle poles, concentration of CPC components at the spindle mid one would automatically favor destruction of cohesin on the short arm. The specific action of each mechanism toward the final outcome of two-step chromosome disjunction remains to be determined.

Subthreshold levels of crossover intermediates are known to induce enrichment of SYP-1 in *cis* onto chromosomes with crossover intermediates along the entire length of the SC, while SYP-1 dissociates from chromosomes without crossover intermediates [45,68]. However, how this crossover intermediate-induced stabilization of SYP-1 relates to partitioning of SYP-1 on short arms is not understood. Our analysis of *dsb-2* mutants has shown that partitioning of both phosphorylated HIM-3 as well as phosphorylated SYP-1 correlates with the number of crossover intermediates. In addition, partitioning of phosphorylated SYP-1 precedes that of unphosphorylated SYP-1, suggesting that phosphorylated SYP-1 has higher affinity toward chromosomes with crossover intermediates. Two possibilities can be considered for the lack of partitioning in nuclei with low numbers of COSA-1 foci: either crossover intermediates generate a signal that must achieve a certain threshold to enable partitioning, or a factor that is normally distributed between 6 chromosomes “spills over” and can no longer enable partitioning when the number of crossover intermediates is low. We expect further genetic studies to illuminate this question.

We found that HIM-3 phosphorylation is dependent on synapsis, raising the possibility of HIM-3 being phosphorylated by PLK-2, whose localization to meiotic chromosomes depends on SYP-1 phosphorylation. However, we still observe phosphorylation of HIM-3 in *plk-2* mutants, and Ser277 and Ser282 are poor matches for a PLK-2 consensus motif. Further investigation is required to identify kinases phosphorylating HIM-3 at these serines as well as at Tyr279. The synapsis-dependent phosphorylation of HIM-3 we observe is the opposite of what has been shown for mouse HORMAD1 phosphorylation at its closure motif, which is detected only on unsynapsed chromosomes [30], suggesting the possibility of a different mode of axis phosphoregulation. Although phosphorylation-mediated regulation of interactions between chromosome axis components may be conserved at the closure motif among HORMA domain proteins, its use is likely to have diverged in order to adapt to different requirements during meiosis between *C. elegans* with holocentric chromosomes and organisms with monocentric chromosomes. Further investigation is needed to determine how post-translational modification of the chromosome axis and SC central elements contribute to their partitioning, function, and disassembly.

## Materials and Methods

### Strains

*C. elegans s*trains were grown with standard procedures [69] at 20°C. Wildtype worms were from the N2 Bristol strain. Mutations, transgenes and balancers used in this study are listed in the **Supplemental Table 2**. For all mutant analyses, we used homozygous mutant progeny of heterozygous parents.

The following point mutants were generated by Clustered Regularly Interspaced Short Palindromic Repeats (CRISPR)-Cas9 gene editing to create point mutations at the endogenous *him-3* locus: *him-3(S277A), (S282A), (AA: S277A S282A) (FA* : *Y279F S282A), (AFA* : *S277A, Y279F, S282A), (S282D)*. In addition, the following mutants were generated in *htp-1::flag* background to enhance HTP-1 visualization and quantification: *(S282A), (AFA: S277A, Y279F, S282A)* and *(S282E)*. For CRISPR-Cas9 gene editing of *him-3* and *syp-1*, gRNAs and synthetic oligonucleotides used as ssDNA homology templates are listed in **Supplemental Table 2**. CRISPR-Cas9 gene editing was done essentially as described in [15] using *dpy-10* as a co-conversion marker [70] with the following modifications: we used crRNA/tracrRNA duplex at 17.5 µM, Cas9 protein at 17.5µM, ssDNA homology template at 6 µM and *dpy-10* ssDNA homology template at 0.5 µM in the injection mix. All the ssDNA homology templates, crRNA and trcrRNA were purchased from Integrated DNA technology (IDT) (**Supplemental Table 2**). The *him-3* and *syp-1(T452A)* homology templates include silent mutations (indicated in lower case in **Supplemental Table 2**) creating XbaI or XspI/MaeI/BfaI cut site, respectively to select for homologous recombination products. Genotyping of *syp-1(T452A)* or *him-3* phospho-mutants was done using the primers listed in **Supplemental Table 2**.

To generate the HTP-1::FLAG fusion, C-terminal tagging of the endogenous *htp-1* locus was performed by injecting the germline with a mix of three plasmids according to [71]. The mix contained: 225 ng/ul of a vector carrying sgRNA htp-1Cterminus2 (TATTTACAGGAACGAGAATA) driven by the U6 promoter, 75 ng/ul of a vector expressing Cas-9 under the *eft-3* promoter, and 150 ng/ul of vector pCFl04 carrying an mCherry reporter and a FLAG tag flanked on both sides by 0.8 kb of sequence homologous to regions upstream and downstream of the Cas9-induced cleavage site at the *htp-1* locus. Selection for homozygous insertion resulted in strain ATG285 *htp-1(fq24[htp-1::FLAG])* IV.

### Phosphoproteomics

Mass spectrometry of phosphoproteins was as described in [16]. To generate the *pph-4*.*1* RNAi plasmid, the 562-bp region of *pph-4*.*1* gene was amplified from *C. elegans* N2 cDNAs using primers 5’-GCTCGT GAA ATC CTA GC-3’ (forward) and 5’-CGA ATA GAT AAC CGGCTC-3’ (reverse) flanked by Not1 and Nco1 sites and cloned into L4440. Peptide counts indicated in Table S1 show pooled counts from all conditions of worms, RNAi, irradiation, or age used in this assay.

### Microscopy, Cytology, Antibodies

For all cytological preparations, we followed protocols described in [72] with the following modification: worm fixation was limited to 2 minutes to improve SC staining. All immunofluorescence was performed on adult worms at 1 day post L4. All the primary antibodies and dilutions used in this study are listed in Supplemental Table 2. Secondary antibodies (Alexa-488, Dylight-594, Dylight-649) were purchased from Jackson ImmunoResearch (PA, USA) and used at 1:500 dilution for all the immunostaining.

Imaging was performed on slides prepared for immunofluorescence on a Deltavision microscope system, using a 100x, 1.4NA PlanSApo objective (Olympus). Images were taken on a Photometrics CCD camera (1024×1024 pixels) at an effective pixel size in the image plane of 64 nanometers. After collecting images, raw 3D data was corrected for lamp flicker and Z-dependent bleaching, then deconvolved using a measured point spread function using the softWoRx suite (GE Healthcare). After deconvolution, wavelengths were offset in the Z direction to correct for chromatic aberration, using multicolor beads as a calibration standard. Quantification of apoptotic nuclei was carried out as described in [58]. For quantification of SYP-1 and HTP-1 localization patterns, oocyte precursor cells from the −4 through −7 diakinesis stage were scored. For quantification of ICP-1 localization pattern, nuclei from −3 through −1 diakinesis stage were scored.

For quantitative analysis of HIM-3 and HIM-3phos intensities (Figure 3), late pachytene nuclei in raw (non-deconvolved) 3D images were classified into “pre” or “post” based on partitioning of HIM-3phos or SYP-2 protein. Ten nuclei of each class were selected per gonad. Within each nucleus, 50 points along the chromosome axis were picked semi-manually in the HIM-3phos channel using the PickPoints feature of the Priism software suite [73]. Points were automatically selected as the maximum intensity pixel within a 5×5×3-pixel box in X, Y, and Z centered on the clicked point, and constrained to ensure points were not duplicated. To compare intensities of pan-HIM-3 on short and long arms, the same nuclei from the “post partitioning” one were used and an additional 50 points were picked in each nucleus in the pan-HIM-3 channel on chromosome stretches where HIM-3-phos points had not been picked previously. To compare intensities between different datasets, points were normalized by dividing all intensity values by the mean intensity of the “pre partitioning” one points. Points were visualized using PlotsOfData [74]. Significance testing was performed in R 3.6.1 (function t.test).

### Embryonic viability quantification

To score embryonic viability of self-fertilized worms, P0 animals at the L4 stage were transferred to new plates every 24 hours. Eggs remaining on the plates 20-24 hours after the transfer were scored as dead eggs. The number of hermaphrodite or male progeny reaching L4/adult stages were scored three days after the transfer. For the *γ*-irradiation experiment, P0 worms at the L4 stage were exposed for 87 minutes at 0.855 Gy/min (total exposure 75Gy) in a Cs-l37 Gammacell 40 Exactor (MDS Nordion) and scored for their embryonic viability as above.

### Fluorescence Polarization (FP) anisotropy peptide assay

Fluorescein (FITC) conjugated peptides for the FP assay were synthesized by either Scrum or Eurofins as shown in **Supplemental Table 2**. HTP-2 proteins were purified and FP assays were conducted as described in [24].

## Acknowledgements

We thank A.F. Dernburg, A. Villeneuve and K. Oegema for antibodies and the past and current members of the Carlton lab for technical assistance. Many nematode strains were provided by the Caenorhabditis Genetics Center, which is funded by the National Institutes of Health National Center for Research Resources. This work was supported by a Japan Society for the Promotion of Science RPD fellowship to A. S-C., Japan Society for the Promotion of Science KAK NHI grants (24687024 Wakate A and l5H04328 Kiban B to P.M.C, and l7Kl5064 Wakate B andl9K06486 Kiban C to A.S-C.), Kyoto university SPIRITS grant to P.M.C., National Institutes of Health grant number R0l-GMl04l4l to K.D.C., and MRC grant MC-A652-5PY60 to E.M-P. No funding bodies had any role in this research.

The authors declare no competing financial interests.

---

**Supplemental Table 1. Counts of phosphopeptides obtained from mass spectroscopy** Details of phosphopeptides detected are shown. For all phosphopeptides detected only once, mass spectrometry spectra were manually examined, and peptides with high confidence assignments are indicated. The MS/MS spectra showing m/z and intensity values for these manually reviewed peptides are shown with accompanying spectrum images in sheets 2 through 4 in the same Excel file.

**Supplemental Figure 1.**
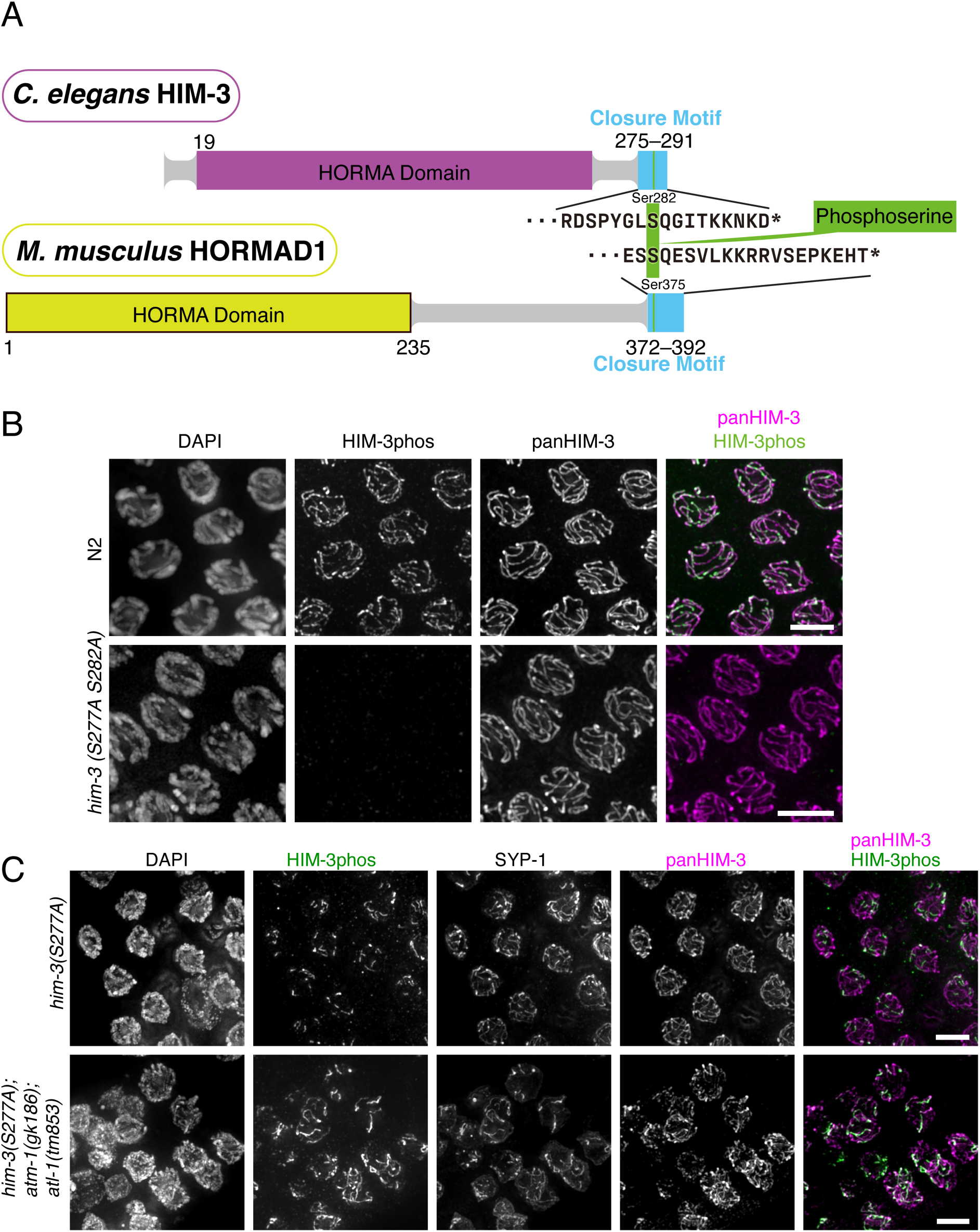
**A**, Diagrams of *C. elegans* HIM-3 (top) and mouse HORMAD1 (bottom), showing the positions of the closure motif and phosphoserine at the C-terminus. **B**, antibodies specific to the phosphorylated closure motif stain the meiotic chromosome axis in wild-type (*top*, green in merged image) but not in S277A,S282A mutants (*bottom*), whereas pan-HIM-3 antibodies stain both (magenta in merged images), verifying the antibody’s specificity to the phosphoepitope. Scalebars, 5µm. **C**, Immunostaining of phospho-HIM-3 (detecting specifically S282 phosphorylation in the *him-3(S277A)* background, green in merged image), SYP-1, and pan-HIM-3 (magenta in merged image) in *him-3(S277A)* (top) and *him-3(S277A); atm-1(gk186); atl-1(tm853)* (bottom) oocyte precursor cells at late pachytene. Although many polyploid nuclei were seen in the *atm-1(gk186); atl-1(tm853)* background, SC structures were partially formed, and HIM-3 within these structures was still phosphorylated.

**Supplemental Figure 2:**
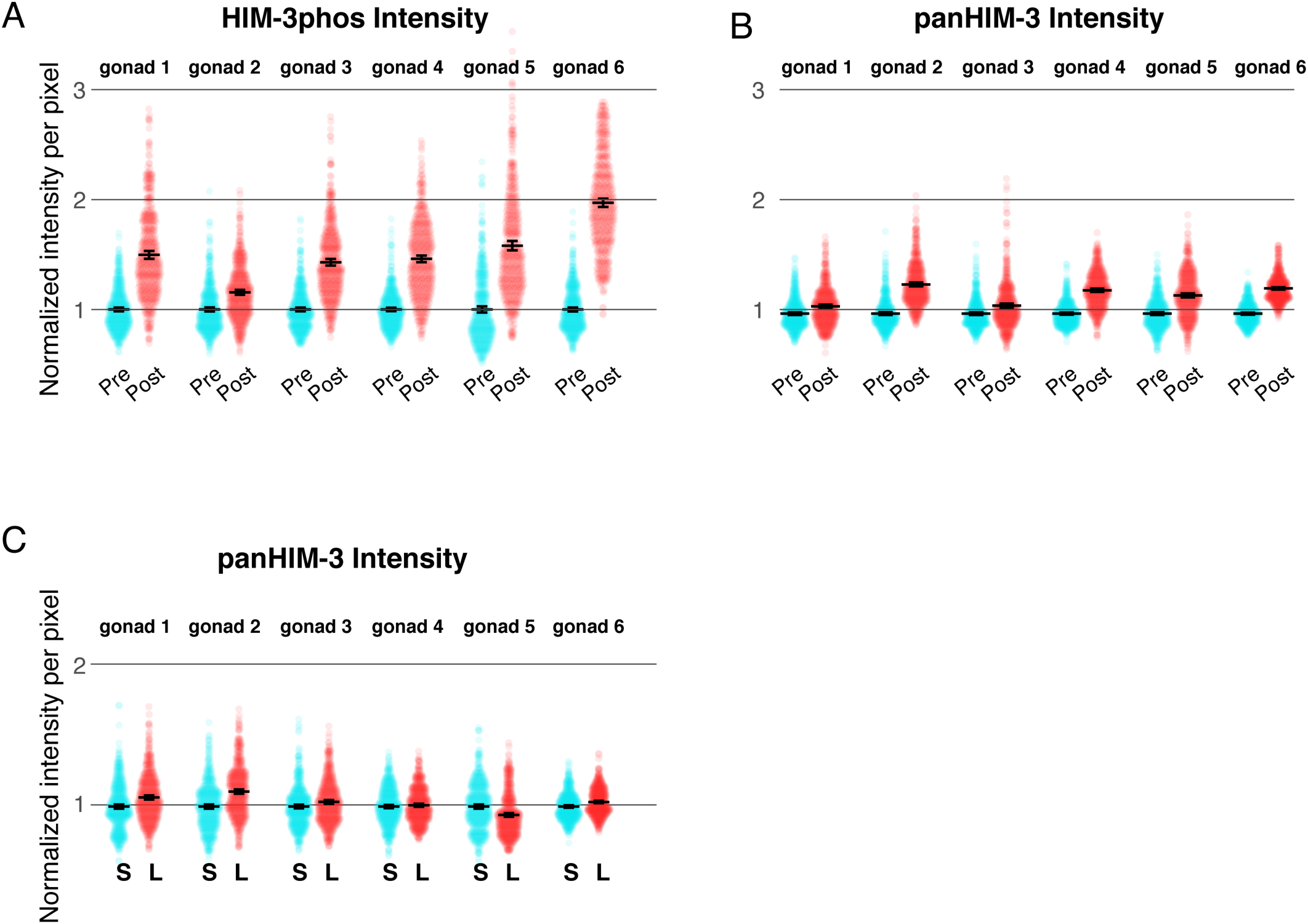
**A, B, C**: individual points plotted for each of the 6 gonads scored for HIM-3 intensity quantitation in **Figure 3**. Comparison of the levels of phosphorylated HIM-3 (A) or panHIM-3 (B) on short arms after partitioning with that of the entire chromosome axis before partitioning within the same gonad. Comparison of the levels of panHIM-3 on short arms with that on long arms after partitioning. Median and 95% confidence intervals are indicated. “Pre” means pre-partitioned nuclei, “post” means post-partitioned nuclei, “S” denotes short arm and “L” denotes long arm.

**Supplemental Figure 3:**
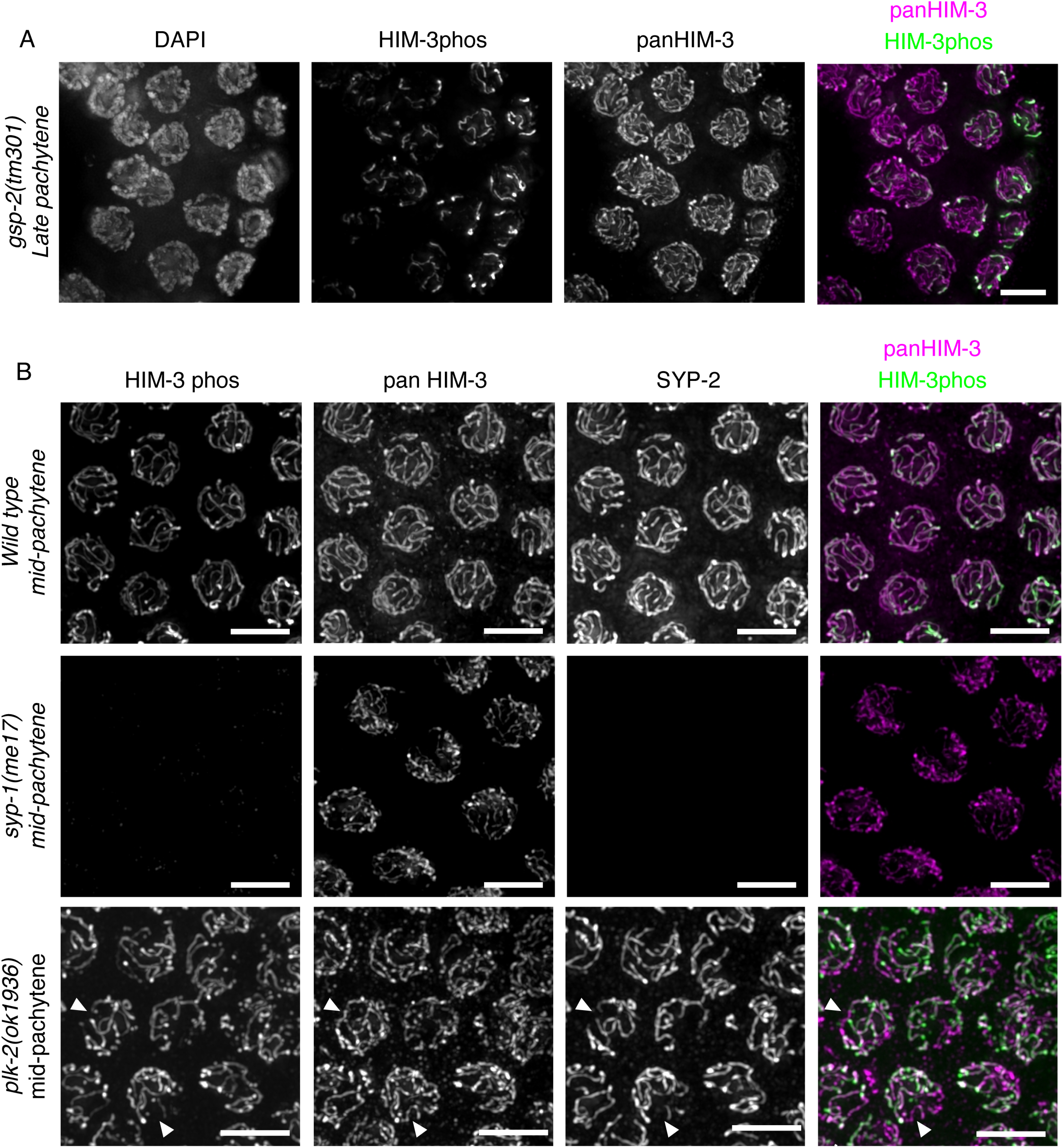
GSP-2 does not dephosphorylate HIM-3, and HIM-3 phosphorylation depends on synapsis. **A**, immunostaining against phospho-HIM-3 (green in merged image) and pan-HIM-3 (magenta in merged image) in a *gsp-2(tm301)* null mutant background. **B**, immunostaining of phospho-HIM-3 (green in merged image), pan-HIM-3 (magenta in merged image) and SYP-2 in the indicated genotypes. In the *plk-2(ok1936)* mutant background (bottom row), arrowheads point to chromosomes that lack synapsis (by a-SYP-2 immunostaining); these same chromosomes also lack phospho-HIM-3 but not pan-HIM-3.

**Supplemental Figure 4.**
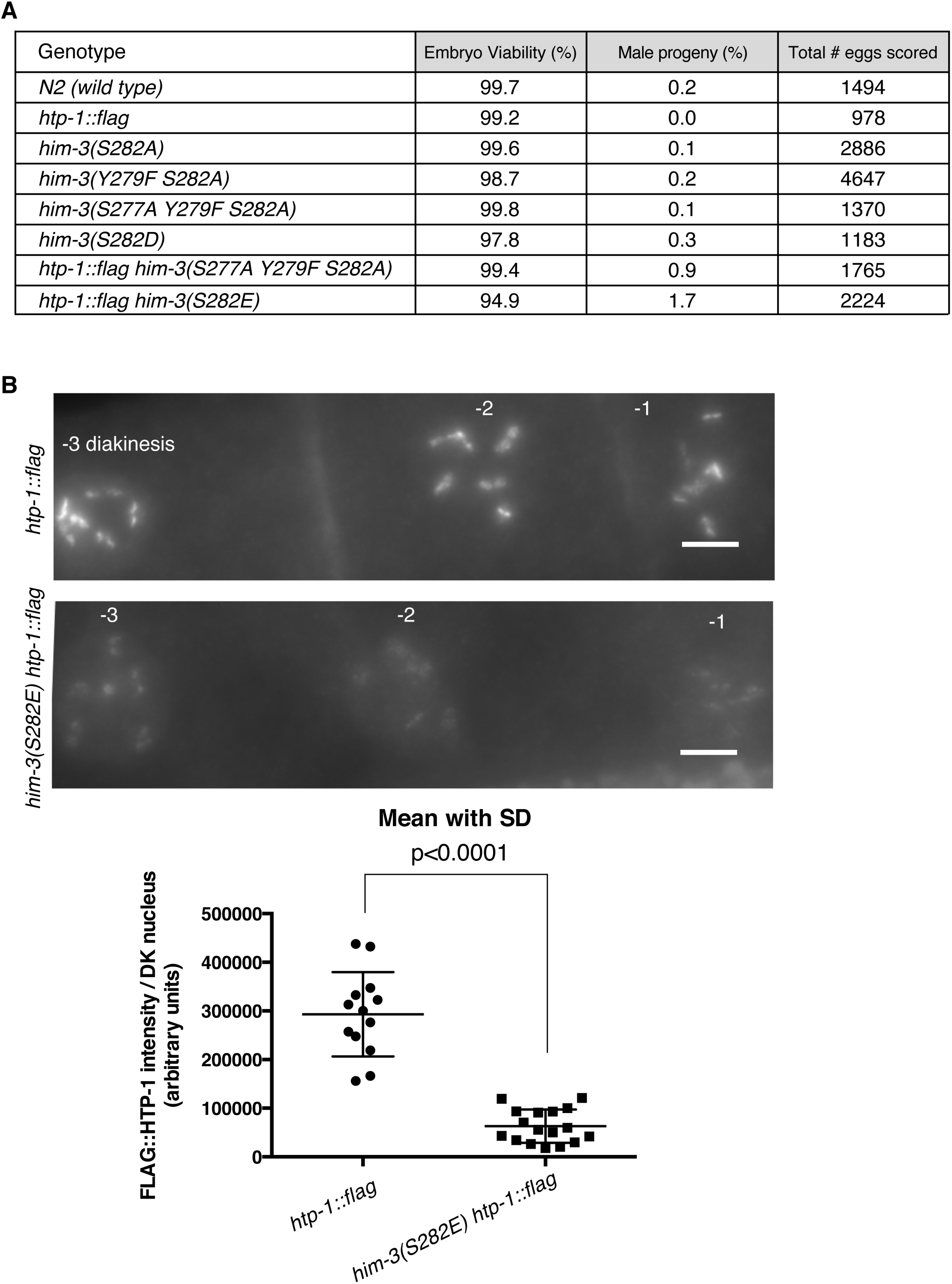
**A**, embryonic viability, male progeny production indicating the rate of *X* chromosome nondisjunction, and total number of scored embryos is shown for the indicated genotypes. **B**, Immunostaining with anti-FLAG epitope on germlines expressing a transgenic HTP-1:FLAG fusion protein in an otherwise wild-type background (top) or a *him-3(S282E)* mutant background (bottom), stained on the same slide, acquired under the same imaging conditions, and displayed with the same scaling. Images shown are summed projections of raw (not deconvolved) image data. Scale bars, 5µm. **C**, quantitation of the intensity of FLAG-HTP-1 immunostaining between the two genotypes in **A**. (Unpaired T-test, p<0.0001)

**Supplemental Figure 5.**
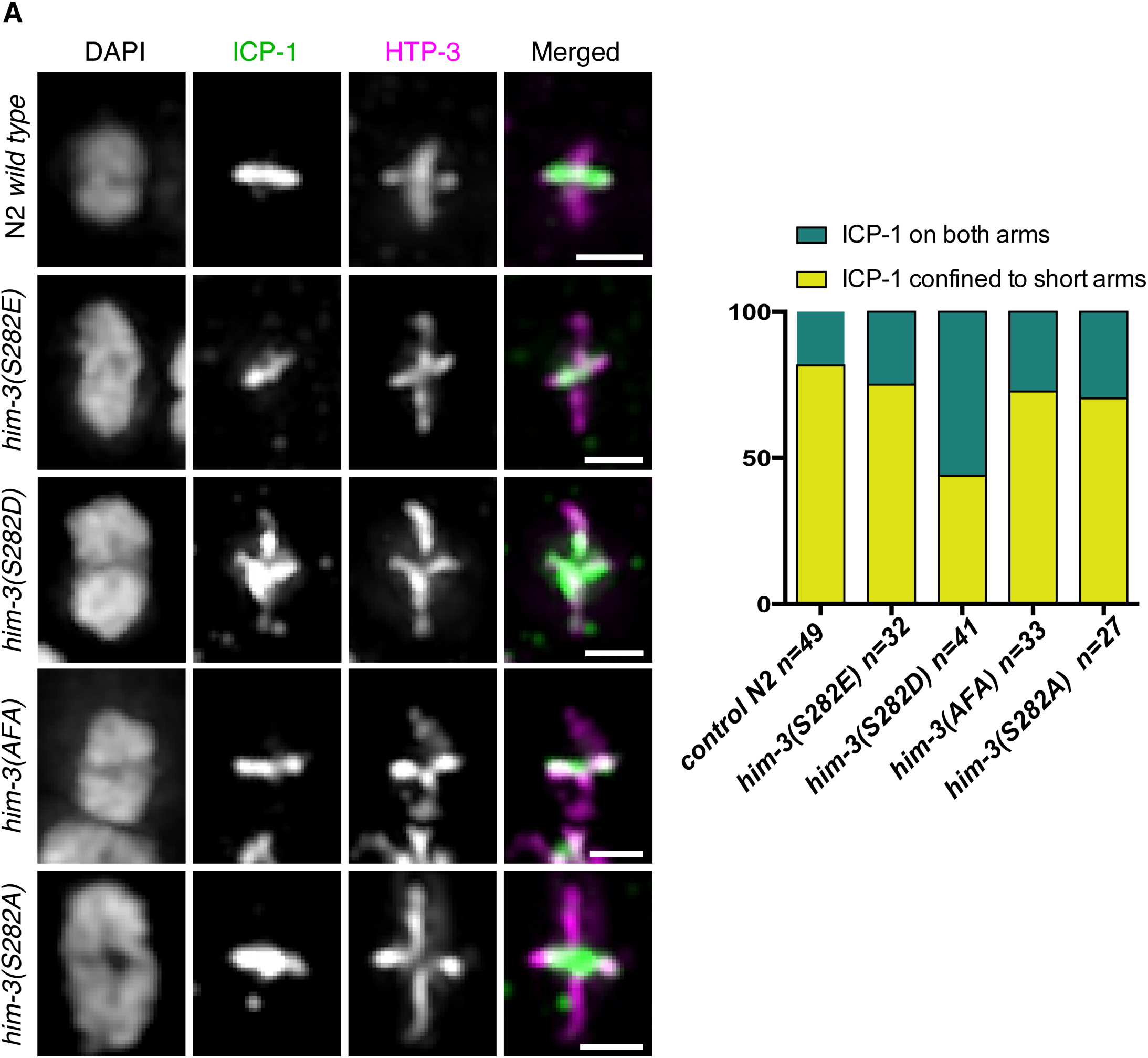
Immunostaining of CPC component INC NP^ICP-1^ (green in merged images) and HTP-3 (magenta in merged images) on individual diakinesis chromosomes in *N2* (wild type), *him-3(S282E) htp-1::flag, him-3(S282D), him-3(AFA) htp-1::flag* and *him-3(S282A) htp-1::flag*. Shown are partial maximum-intensity projections of single chromosomes oriented so the long arms are vertical. Quantification of ICP-1 localization classes (confined or unconfined) is shown at right.

**Supplemental Figure 6.**
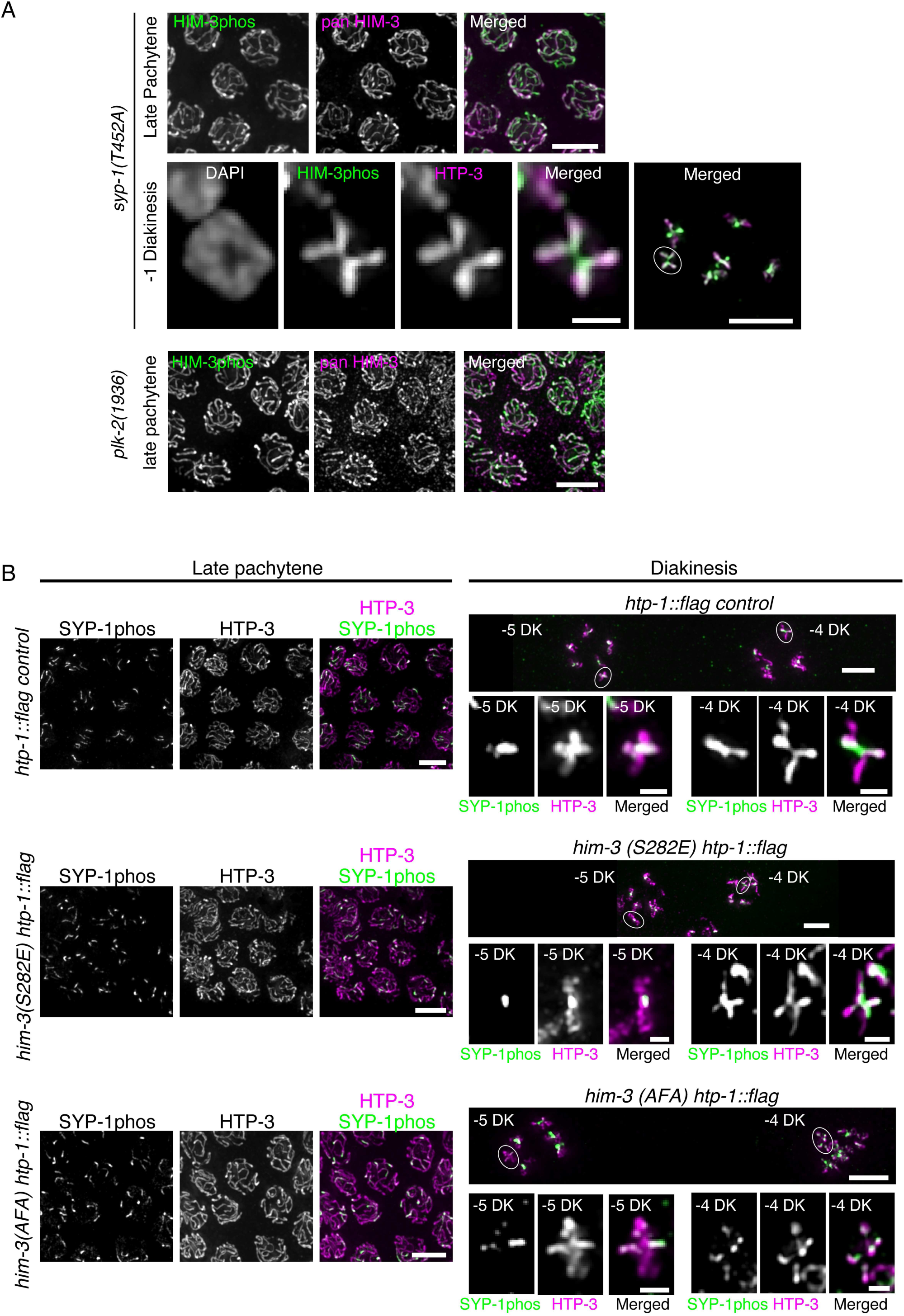
**A**, Immunostaining of phospho-HIM-3 (green in merged images) and pan-HIM-3 (magenta in merged images) in the indicated genotypes showing phospho-HIM-3 failing to partition to the short arm. Partial Z projections are shown for all images. **B**, Immunostaining of phospho-SYP-1 (green in merged images) and HTP-3 (magenta in merged images) in the indicated genotypes. For both A and B, individual diakinesis chromosomes displayed are circled in the overview images. In *him-3(S282E)* mutants, although SYP-1phos staining is confined normally to short arms from late pachytene up to −5 diakinesis, SYP-1phos signals reappearing on long arms are occasionally found specifically in −4 and −3 diakinesis nuclei. Scalebars: individual diakinesis chromosomes, 1µm; all others 5µm.

**Supplemental Figure 7.**
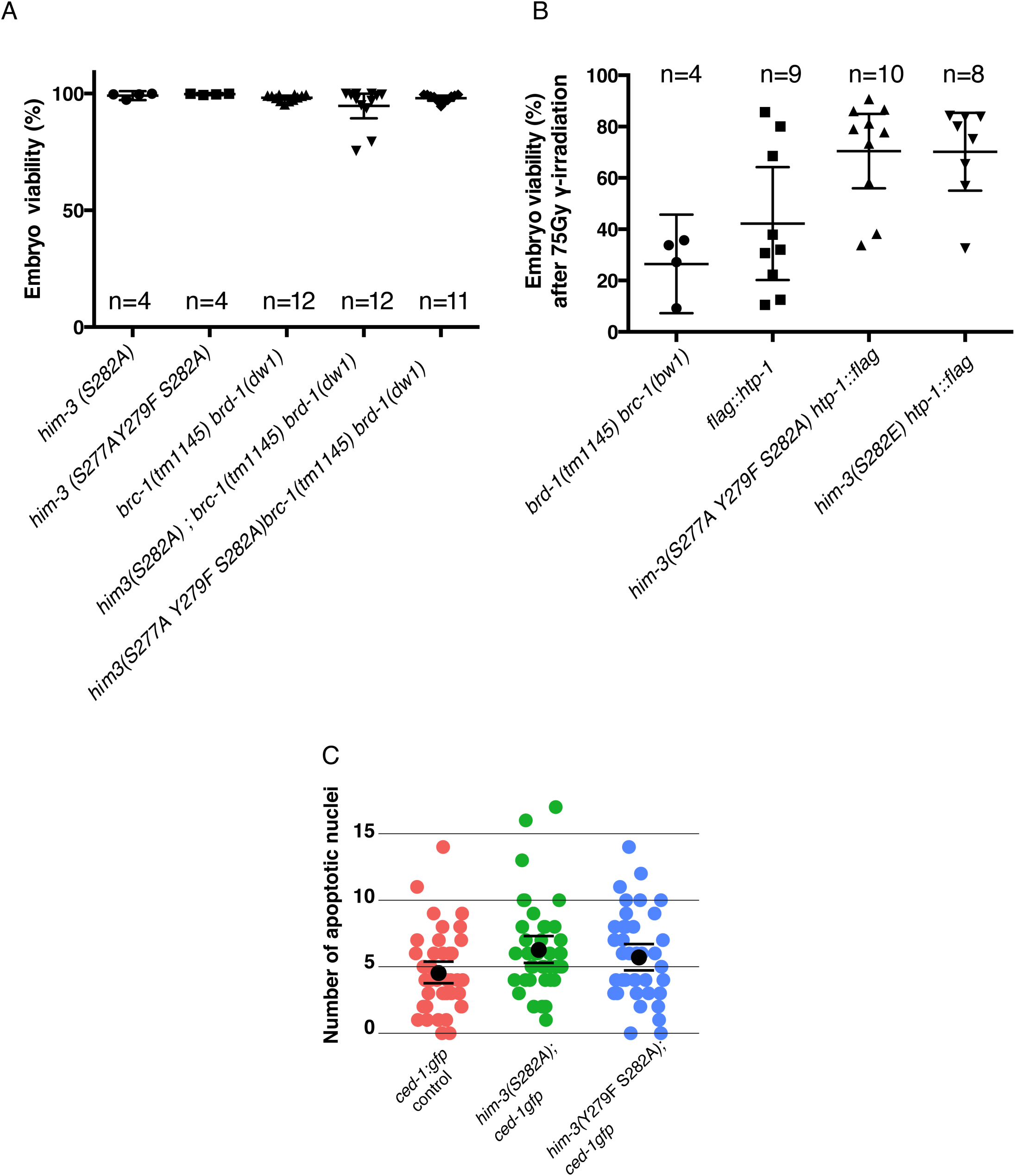
**A**, Percentage of embryonic viability among the self-progeny of worms with the indicated genotypes. For each column, n indicates the number of maternal worms. Total number of eggs scored are : *him-3(S282A)*:ll85, *him-3(AFA)*: l370, *brc-1(tm1145) brd-1(dw1)*: 3013, *him-3(S282A); brc-1(tm1145) brd-1(dw1)*: 2l78, *him-3(AFA); brc-1 brd-1*: 2443 eggs. Mean with 95% confidence intervals is indicated in the graph. The *him-3* non-phosphorylatable mutations did not lower embryonic viability when sister-chromatid-mediated homologous recombination is prevented in the *brc-1(tm1145) brd-1(dw1)* background. **B**, Percentage of embryonic viability among the self-progeny of P0 worms after 75 gray gamma-irradiation at L4 stage. The *brc-1(tm1145) brd-1(dw1)* mutant was used as a positive control, which shows reduced embryonic viability upon gamma irradiation. Total number of eggs scored are : *brc-1(tm1145) brd-1(dw1):* 646, *htp-1::flag* :1178, *him-3(AFA) htp-1::flag*: l896, *him-3(S282E) htp-1::flag:* 1233 eggs. Mean with 95% confidence intervals is indicated in the graph. The *him-3* phospho-mutations did not lower embryonic viability upon exogenous DNA damages. **C**. scoring of apoptotic nuclei in *him-3* phospho-mutant backgrounds. The number of C D-l:GFP-positive nuclei for each condition is plotted with mean and 95% confidence intervals. The *him-3* non-phosphorylatable mutations did not increase the number of apoptotic nuclei.

